# Processing of forked DNA activates a helicase-nuclease immune system

**DOI:** 10.64898/2026.05.13.724313

**Authors:** Luuk Loeff, Christelle Chanez, Martin Jinek

## Abstract

Innate immune systems detect molecular signatures of infection to initiate antiviral defence^1–3^, yet the identity of pathogen-associated signals that distinguish phage from host nucleic acids remains incompletely understood. While recent work has shown that nucleic acid structures can act as triggers for bacterial defense systems^4–7^, how these structural signals are coupled with immune activation remain unclear. Here we show that forked DNA structures activate a helicase-nuclease immune complex in type III Druantia through a processing-dependent mechanism. Using cryo-electron microscopy and biochemical reconstitution, we find that the exonuclease DruH processes 3′ DNA termini to generate 5′ overhangs that recruit and activate the helicase-nuclease DruE at duplex-single-stranded DNA junctions. Structural analysis of the DruE-DruH complex reveals how substrate-dependent assembly remodels an autoinhibited helicase dimer into an active DNA degradation complex. Functional assays demonstrate that coordinated nuclease and helicase activities enable efficient degradation of forked DNA substrates and mediate phage defense without detectable host toxicity. Together, our findings define a mechanism in which enzymatic processing of replication-associated DNA structures licenses immune activation, providing a framework for how nucleic acid architecture is coupled to effector activation in bacterial immunity.

## Introduction

Prokaryotic organisms are estimated to be outnumbered tenfold by their viruses (bacteriophages), resulting in 10^25^ microbial infections per second^8^. This constant host–parasite conflict has driven the evolution of a diverse array of defense systems that protect against viral invasion. Genes encoding prokaryotic defense systems are often organized into clusters, commonly referred to as defense islands^9^. This organization has enabled systematic discovery of novel defense systems through computational and functional approaches^10–12^, uncovering more than two hundred defense systems with diverse enzymatic activities^13–17^. Although the roles of many systems in antiviral defense have been established, the molecular features they recognize and the mechanisms by which they are activated remain incompletely understood.

Emerging evidence suggests that prokaryotic immune systems detect nucleic acid structures to distinguish phage from host DNA, including DNA ends and replication-associated intermediates^4–7^. Druantia is a widespread prokaryotic host defense system that confers broad phage immunity^13,18^, and is defined by the presence of the druE signature gene, which encodes a ∼2000 amino acid ATPase/helicase protein^13^. Type III Druantia systems comprise DruE and DruH, a large accessory protein with unknown function. Due to the size of the system and the lack of recognizable domains in the accessory genes, Druantia is thought to function as a multi-component molecular machine. However, the molecular mechanism by which Druantia detects phage infection and activates its effector functions has remained unresolved.

In this study, we used an integrative approach combining single-particle cryo-electron microscopy (cryo-EM), biochemistry and phage infection assays to define the mechanism of type III Druantia. Our biochemical and structural analysis shows that DruE is an autoinhibited dimer, whereas the multidomain protein DruH functions as a 3’ to 5’ exonuclease. A cryo-EM structure of the DruE-DruH holocomplex and complementary biochemical assays reveal that DruH-dependent recognition of forked DNA triggers DruE dimer disassembly and loading onto the 3′ strand at duplex–ssDNA junctions. Complementary unwinding assays demonstrate that DruE preferentially initiates on DNA substrates containing 5’ single-stranded overhangs. These findings establish a mechanism in which enzymatic processing of replication-associated DNA structures licenses immune activation, enabling selective targeting of phage DNA while avoiding host toxicity.

## Results

### Phylogenetic analysis of DruE reveals four distinct Druantia subtypes

To characterize the diversity of Druantia systems, we performed a phylogenetic analysis of 566 Druantia loci using the conserved helicase signature gene *druE* as a phylogenetic marker. The DruE orthologs segregated into four distinct clades (**Figure 1A**), hereafter referred to as Type I to IV, consistent with previously described Druantia types^13,19,20^. Among these, Type III systems are the most abundant encompassing 46% of the Druantia loci, whereas type I, II and IV encompass 14%, 22% and 18% of loci, respectively (**Extended data Fig. 1A**). Together, these results define the phylogenetic landscape of Druantia systems and motivate further structural and mechanistic characterization of Type III as the most prevalent Druantia system.

**Figure 1:**
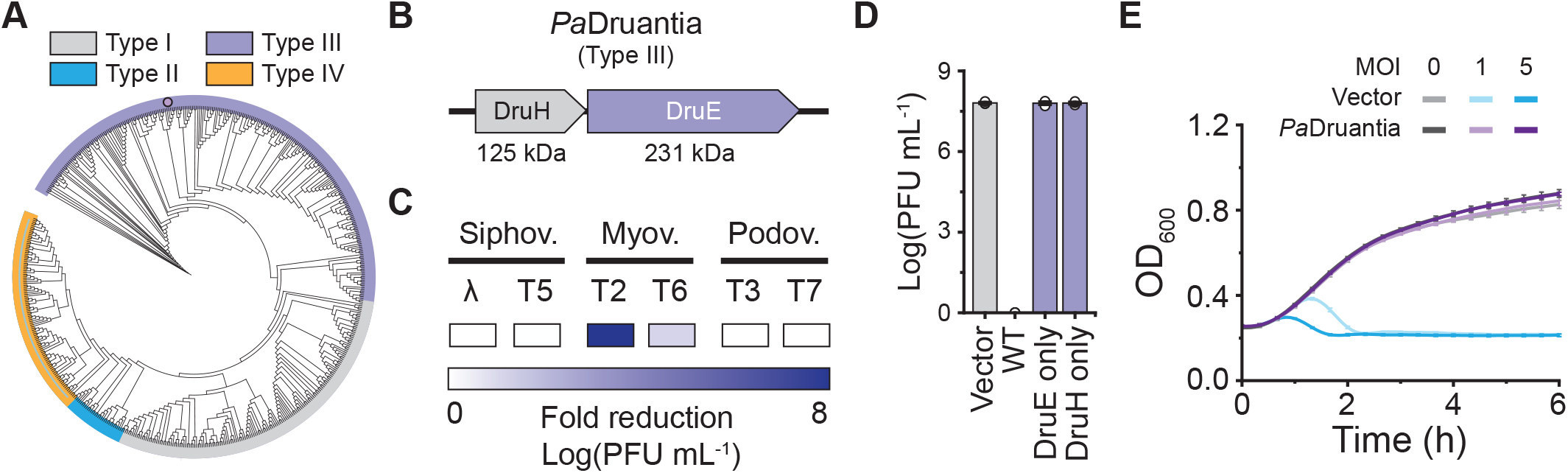
Phylogeny and phage defense by type III Druantia. **(A)** Phylogenic classification of Druantia systems based on 566 DruE orthologs into four distinct subtypes. **(B)** Genomic architecture of Type III Druantia in Pseudomonas aeruginosa BL21. DruH (grey) and DruE (purple). **(C)** Phage defense phenotype of *E. coli* cells expressing type III *Pa*Druantia against six *E. coli* phages. Color intensity represents fold reduction in the efficiency of plating (EOP), also see Extended data Fig. 1C. **(D)** Plaque assays of T2 phage infecting *E. coli* strains expressing wild-type (WT) PaDruantia, PaDruH alone, PaDruE alone or a vector-only control. Data represents mean PFU mL^-1^ ± SEM of three independent replicates (n=3). **(E)** Growth curves for *E. coli* cells harboring the PaDruantia system or an empty vector, infected with the T2 phage at a multiplicity of infection (MOI) of 0, 1 or 5. Data points represent mean ± SEM of three biological replicates (n=3).

### Type III Druantia provides phage immunity

To elucidate the molecular mechanism of Type III Druantia, we focused on the *Pseudomonas aeruginosa* Druantia system (PaDruantia, **Figure 1B**). As prokaryotic defense systems can cause growth defects or toxicity when ectopically expressed in a heterologous host, we assessed whether Type III Druantia expression is intrinsically toxic in *E. coli* by expressing PaDruantia cells using an arabinose-inducible promoter. Cells expressing either PaDruantia wild-type, DruH alone or DruE alone, grew at similar rates compared to cells harboring an empty expression vector (**Extended data Fig. 1B**), indicating that PaDruantia activity does not result in nonspecific DNA damage or growth inhibition in the host cell. These observations indicate that Type III Druantia does not function as a canonical toxin–antitoxin system, as expression of individual components does not impair host growth in the absence of phage challenge.

Next, we challenged *E. coli* harboring PaDruantia with a set of six of *E. coli* phages that belong to the three major families of tailed dsDNA phages (*Sipho-, Myo-*, and *Podoviridae*). PaDruantia provided a strong defense response against *Myoviridae* bacteriophages T2 and T6, resulting in >1×10^7^-fold and >2×10^4^-fold reductions in the efficiency of plating (EOP), as compared to the vector-only control (**Figure 1C & Extended data Fig. 1C**), respectively. By contrast, no defense was observed for the phages belonging to the *Siphoviridae* and *Podoviridae* families. Subsequently, we challenged *E. coli* strains harboring either DruE or DruH with T2 phages and observed no defense phenotype, indicating that simultaneous presence of DruE and DruH is essential for phage defense (**Figure 1D**). Collectively, these observations indicate that Type III Druantia provides defense against a subset of Myoviridae phages.

Over 70% of prokaryotic gene defense systems characterized to date mediate immunity by causing programmed cell death to abort phage infection^21,22^, we assessed whether PaDruantia provides phage defense through an abortive infection mechanism. When cells lacking PaDruantia were challenged with T2 phage at a low and high multiplicity of infection (MOI) in liquid culture, cell cultures collapsed due to the lytic replication cycle of T2. By contrast, cells harboring PaDruantia resisted infection even at a high MOI and grew at the same rate as the uninfected control **(Figure 1E**), indicating that Druantia does not elicit programmed cell death upon phage infection. This suggests that Druantia instead provides immunity through a mechanism that inhibits phage propagation and implies that Druantia systems are capable of self-versus-non-self-discrimination.

### DruH functions as a DNA targeting exonuclease

To understand the defense mechanism of Type III Druantia, we sought to obtain structural insights into the DruH protein. Recombinant expression and purification of DruH yielded a monodisperse protein sample, as indicated by size exclusion chromatography (**Extended data Fig. 1D**). DruH was subjected to structural analysis by single-particle cryo-EM, which yielded a reconstruction with a nominal resolution of 3.0 Å (**Figure 2A and Extended data Fig. 2A-D**). The cryo-EM structure of DruH reveals a monomer comprised of an N-terminal alpha helical domain, six Immunoglobin-like (Ig-like) domains and a C-terminal alpha helical domain (**Figure 2B**). The N- and C-terminal domains interact through an extensive electrostatic interface that spans a surface area of 13774 Å^2^.

**Figure 2:**
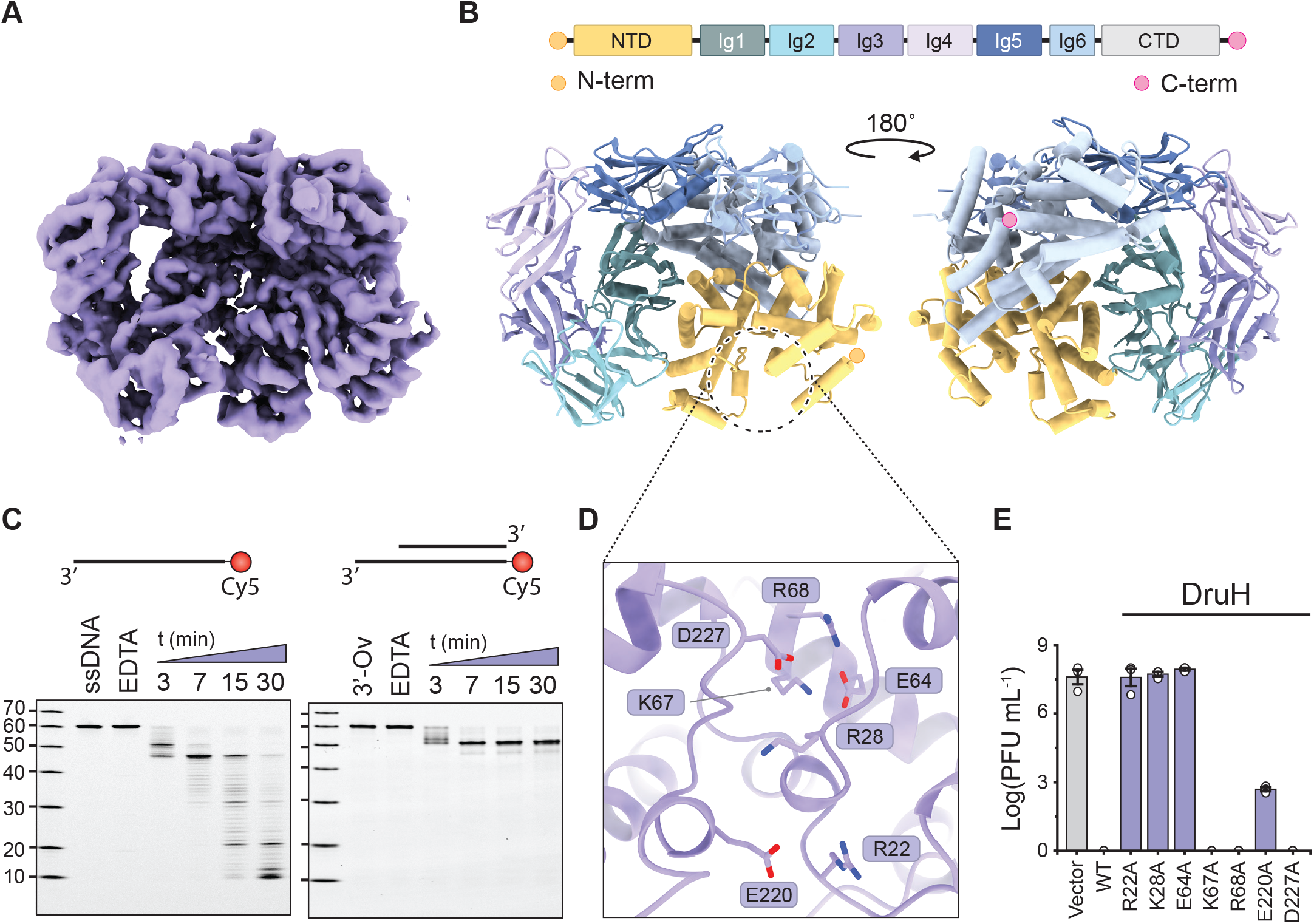
DruH is a multidomain nuclease required for Druantia-mediated defense. **(A)** Cryo-EM density map of the DruH protein. **(B)** Structural model of DruH, shown in cartoon representation. N- and C-termini are indicated by orange and pink circles, respectively. **(C)** *In vitro* nuclease activity assays with DruH using fluorescently labelled oligonucleotides. Cleavage products were resolved by denaturing polyacrylamide gel electrophoresis (PAGE), also see **Extended data Fig. 3D-F. (D)** Close-up view of the putative active site region of PaDruH showing highly conserved residues, also see **Extended data Fig. 3G-H. (E)** Plaque assays of T2 phage infecting *E. coli* strains expressing wild-type (WT) PaDruantia, PaDruH mutants, or a vector-only control. Data represents mean PFU mL^-1^ ± SEM of three independent replicates (n=3).

Structure-based homology searches of DruH identified high-confidence similarities to human Ig-like domains found in intercellular adhesion molecules such as ICAM and NCAM (**Extended data Fig. 3A**). However, while these hits indicate structural conservation, they do not provide insight into the molecular function of DruH. Consistently, sequence-based (InterPro) searches did not yield any significant matches. Given the predicted helicase function of the signature gene *druE* we hypothesized that Type III Druantia systems directly target phage-derived nucleic acids. To investigate whether DruH mediates nucleic acid recognition or processing, we examined AlphaFold3 structural predictions of DruH in the presence of various nucleic acid structures. When DruH was co-folded with a deoxyribonucleic acid (DNA) substrate containing 3’ overhang, the single-stranded DNA (ssDNA) overhang was threaded through DruH. In contrast, the 5’ overhang substrate was not accommodated within DruH and remained bound at the protein surface (**Extended data Fig. 3B-C**).

To test whether these structural predictions are reflected in the biochemical activity of DruH, we tested whether DruH possessed nuclease activity in vitro using fluorophore-labeled synthetic oligonucleotide substrates. Incubation of DruH with ssDNA in the presence of Mg^2^+ led to complete degradation of the oligonucleotide, while substrates containing a 3′ overhang were progressively degraded until the dsDNA junction (**Figure 2C**). In contrast, blunt dsDNA, single-stranded ribonucleic acid (ssRNA) or DNA substrates containing a 5′ overhang did undergo detectable degradation in the presence of DruH and Mg2+ (**Extended data Fig. 3D-F**). These results suggest that DruH acts as a DNA-targeting exonuclease with 3′–5′ polarity. Structural predictions placed the DNA substrate within a cleft in the N-terminal α-helical domain of DruH, which is lined by highly conserved residues forming a putative catalytic site (**Extended data Fig. 3G-H**). Alanine substitutions of Arg22^DruH^, Lys28^DruH^, and Glu64^DruH^ abolished Druantia-mediated defense against phage T2, whereas mutation of Glu220^DruH^ resulted in a partial reduction of immunity in vivo (**Figure 2E**), supporting a functional role for this conserved surface in DruH activity. Together, these results establish DruH as a DNA-specific 3′-5′ exonuclease that selectively targets exposed single-stranded DNA at 3′ termini and DNA junctions, suggesting a role in processing structured DNA substrates generated during phage infection.

### DruE functions as a helicase-coupled nuclease

To gain mechanistic insights into the activity of the Druantia signature protein DruE, we recombinantly expressed, purified (**Extended data Fig. 1E**), and determined its the structure by single-particle cryo-EM, obtaining a reconstruction with a nominal resolution of 3.1 Å (**Figure 3A & Extended data Fig. 4A-D**). The structure reveals a pseudo-C2 symmetric homodimeric assembly in which the individual protomers interact via an extensive electrostatic interface that spans a surface area of ∼1907 Å^2^ (**Figure 3B**). Despite the overall symmetry, the reconstruction showed asymmetry in terms of local resolution likely due to conformational flexibility. As a result, only one protomer was sufficiently well resolved to enable atomic model building (**Figure 3A-B & Extended data Fig. 4C**). The domain organization of the DruE N-terminal helicase closely resembles that of *Bacillus subtilis* MrfA, a processive 3’-5’ translocating SF2 DNA helicase involved in DNA repair that uses a distinctive skipping rope translocation mechanism (**Extended data Fig. 5A**)^**23**^.

**Figure 3:**
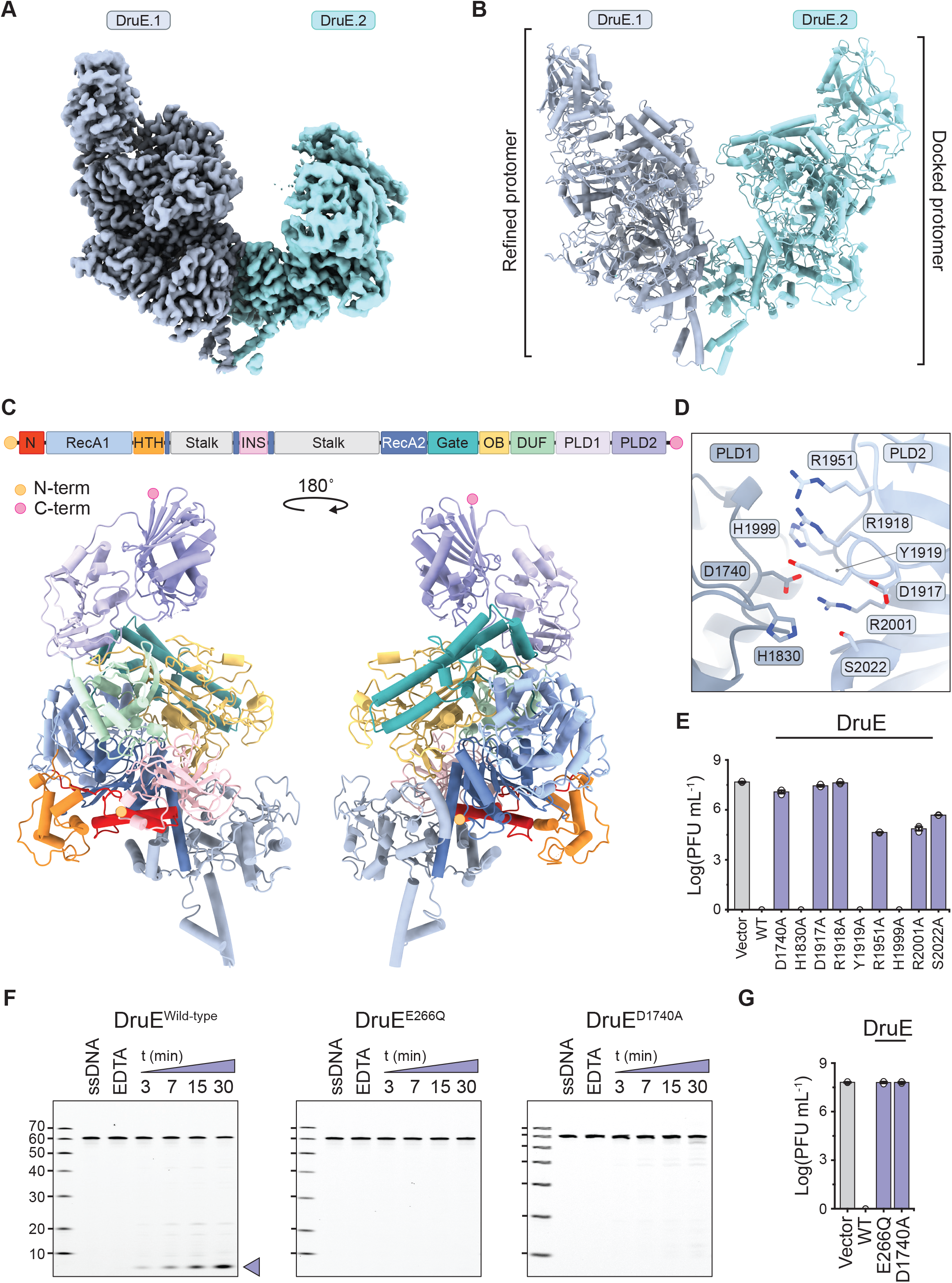
The helicase-nuclease DruE froms an autoinhibited dimer. **(A)** Cryo-EM density map of the dimeric DruE complex. **(B)** Structural model of dimeric DruE, shown in cartoon representation and in the same orientation as in A. **(C)** Structural model of a DruE protomer (bottom) shown alongside domain organisation (top). N- and C-termini are indicated by orange and pink circles, respectively. **(D)** Close-up view of the PLD nuclease active site of DruE showing its catalytic residues. **(E)** Plaque assays of T2 phage infecting *E. coli* strains expressing wild-type (WT) PaDruantia, PaDruE catalytic mutants, or a vector-only control. Data represents mean PFU mL^-1^ ± SEM of three independent replicates (n=3). **(F)** *In vitro* nuclease activity assays with WT DruE, or DruE proteins carrying inactivating mutations in the helicase (E266Q) or nuclease (D1740A) domains. using fluorescently labelled ssDNA oligonucleotides. Cleavage products were resolved by denaturing polyacrylamide gel electrophoresis (PAGE) **(G)** Plaque assays of T2 phage infecting *E. coli* strains expressing wild-type (WT) PaDruantia, PaDruE helicase (E266Q) or nuclease (D1740A) mutants, or a vector-only control. Data represents mean PFU mL^-1^ ± SEM of three independent replicates (n=3).

The helicase core of DruE is composed of two canonical RecA-like helicase domains (RecA1 and RecA2), which together form an ATPase active site located at their interface (**Figure 3C**). Two additional domains, hereafter referred to as Insertion (INS) and Stalk, are embedded within the RecA2 domain DruE. The Stalk domain mediates dimerization and contains a long alpha helix that protrudes from the RecA2 core, whereas the INS domain is positioned at the nucleic acid entrance to the helicase (**Figure 3C**). Apart from its helicase core, DruE contains six additional domains: N-terminal (N), a helix-turn-helix, Gate, DUF1998 (DUF), oligonucleotide binding (OB) and a Phospholipase D (PLD) domains. Of these, the N, DUF1998 and OB domains are found in MrfA (**Figure 3C**)^23^. DruE additionally contains several Cys-rich zinc-binding motifs, which may contribute to the structural integrity of the protein. This overall domain architecture is consistent with the recently proposed type III-A classification of Druantia systems^24^.

Superposition of the DruE dimer with DNA-bound MrfA (PDB: 6ZNP) reveals that the Gate domain of DruE blocks the helicase exit channel, resulting in severe clashes with modeled DNA, indicating that productive DNA engagement requires substantial conformational rearrangements (**Extended data Fig. 5B-C**). To test DNA binding, we used electrophoretic mobility shift assays. DruE was capable of binding partial duplex substrates containing either 3′ or 5′ single-stranded DNA overhangs (**Extended data Fig. 5D-E**), whereas blunt-ended dsDNA failed to form a discrete complex (**Extended data Fig. 5F**). Interestingly, DNA binding was accompanied by a shift toward monomeric DruE, indicating that dimerization impedes productive DNA engagement (**Extended data Fig. 5D-E**). This behavior is reminiscent of the *Vibrio cholerae* DdmDE plasmid defense system, in which the autoinhibited helicase nuclease DdmD dimer dissociates upon activation by the DNA bound prokaryotic Argonaute DdmE^25–27.^

The C-terminal region of DruE projects over the helicase core and contains two PLD-like domains that closely resemble Human Microtubule Interacting and Trafficking Domain Containing 1 (MITD1) protein (**Extended data Fig. 5G**). As the PLD superfamily includes nuclease enzymes^28,29^, the presence of PLD domains in DruE suggests that it may have nuclease activity. A conserved pocket lined with residues Asp1740^DruE^, His1830^DruE^, Asp1917^DruE^, Arg1918^DruE^, Tyr1919^DruE^, Arg1951^DruE^, His1999^DruE^, Arg2001^DruE^, Ser2022^DruE^ forms a putative nuclease active site (**Figure 3D & Extended data Fig. 5H-I**). Accordingly, alanine substitutions of Asp1740^DruE^, Asp1917^DruE^, Arg1918^DruE^ abolished Druantia-mediated phage defense against T2 phage *in vivo*, while subsitution of Arg1951^DruE^, Arg2001^DruE^, Ser2022^DruE^ resulted in an intermediate reduction in anti-phage activity (**Figure 3E**). Subsequently, we analyzed DruE-mediated nuclease activity *in vitro* using a fluorophore-labeled synthetic single stranded oligonucleotide substrate. Incubation of wild-type DruE with ssDNA, Mg^2+^ and ATP resulted in the accumulation of fully digested nucleic acid products, whereas mutation of the helicase (E266Q, designed to impair ATP hydrolysis, **Extended data Fig. 5J**) or nuclease domains (D1740A) abolished DNA degradation *in vitro* and phage defense in *E. coli* (**Figure 3F-G**). Collectively, these results indicate that DruE functions as an ATP-driven nuclease in Druantia-based immunity, whose activity is inhibited by dimerization.

### DruE-DruH complex formation primes DruE for DNA degradation

We next focused on the mechanism underpinning phage sensing and self-versus-non-self discrimination by type III Druantia. Given the strong activity of PaDruantia against T-even phages of the *Myoviridae* family (**Figure 1C**) and the recently reported escape of phages encoding a methyltransferase from type I Druantia immunity^30^, we initially hypothesized that Type III Druantia may specifically recognize the glucosylation of 5-hydroxymethylated cytosines found in the DNA of T-even phages^31,32^. However, glucosylated genomic DNA from T2 or T6 phages did not exhibit substantial degradation in the presence of DruH and DruE *in vitro*, indicating type III Druantia does not specifically recognize glycosylated dsDNA substrates (**Extended data Fig. 6A-B**). This observation contrasts with robust cleavage of oligonucleotide substrates by DruH and DruE (**Figure 2C & 3F**), suggesting that Druantia Type III may instead target DNA intermediates that occur during phage replication.

Given that DruE and DruH are both nucleases, we hypothesized that they form a complex upon engagement with a cognate DNA substrate relevant to Druantia activity. To test this, we performed affinity co-precipitation experiments using matrix-immobilized DruH in the presence of defined synthetic DNA substrates. When DruH was incubated with a forked DNA substrate, we observed efficient DruE co-precipitation (**Figure 4A-B**). By contrast, DruE was not co-precipitated by DruH alone or in the presence of ssDNA, dsDNA or a dsDNA replication bubble mimic substrate containing internal mismatched sequences (**Figure 4A**). To validate the functional relevance of this observation, we reconstituted DNA degradation activity of the type III Druantia system *in vitro*. Upon incubation of a double fluorescently labelled forked DNA substrate (**Figure 4B**) with DruE and DruH, both strands of the substrate were degraded (**Figure 4C**). By contrast, mutation of the helicase (E266Q, designed to impair ATP hydrolysis, **Extended data Fig. 5J**) abolished DNA degradation while a nuclease mutant (D1740A) exhibited only weak residual degradation (**Figure 4C**). Notably, DruE alone exhibited detectable nuclease activity, albeit at substantially reduced efficiency, suggesting that DruH enhances DNA degradation, potentially by stabilizing the degradation-competent complex (**Figure 4C**).

**Figure 4:**
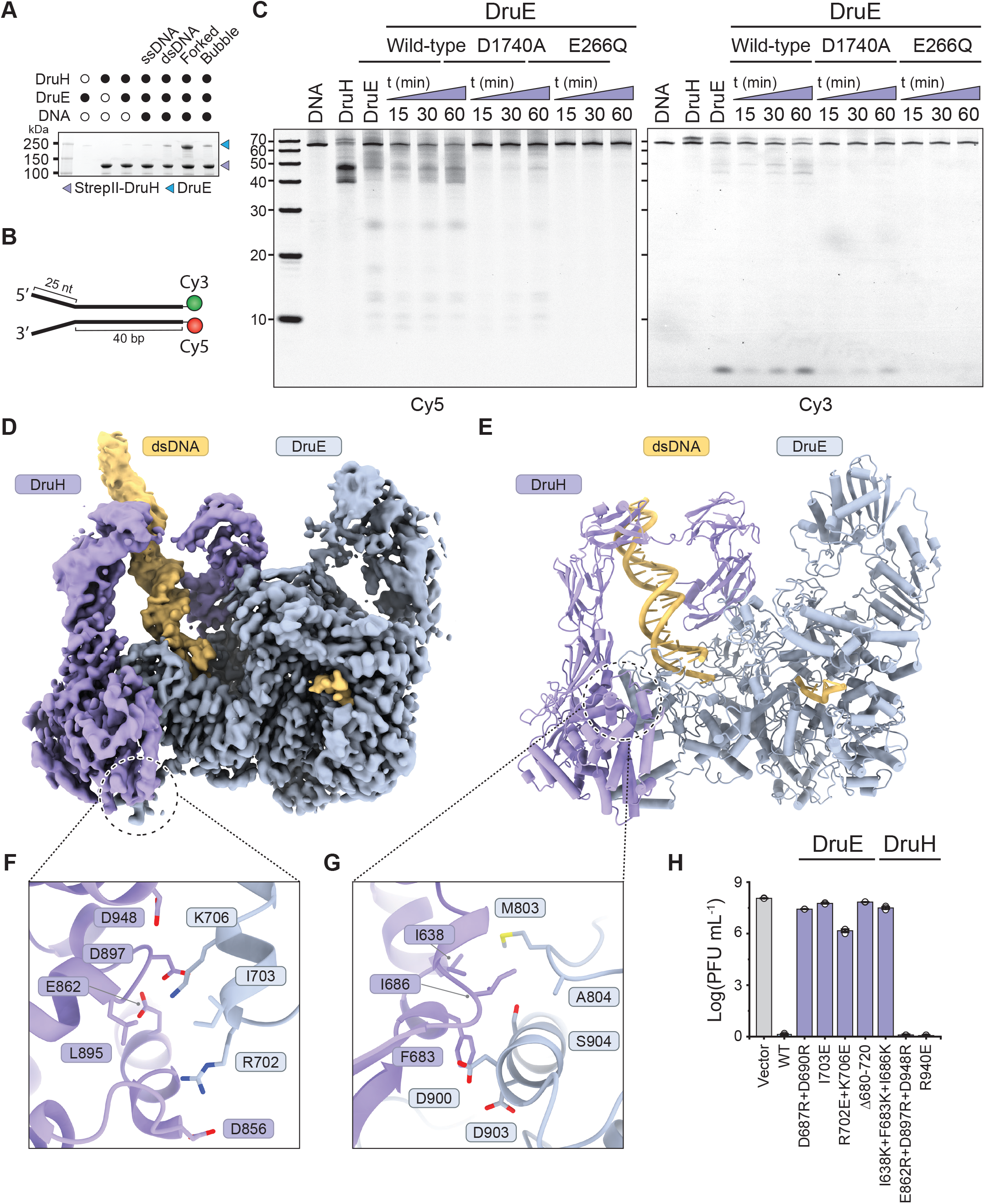
DNA-dependent assembly and structure of the DruE-DruH complex. **(A)** *In vitro* pulldown experiment examining DNA dependent complex formation between recombinant DruE and StrepII-tagged DruH in the presence of defined DNA substrates, including a ssDNA, dsDNA, forked DNA substrate and a dsDNA replication-bubble mimic containing a mismatched central region (bubble). **(B)** Schematic diagram of the forked DNA substrate used for the *in vitro* reconstitution assay and cryo-EM complex assembly. **(C)** *In vitro* nuclease activity assay with WT Druantia or DruE proteins carrying inactivating mutations in the helicase (E266Q) or nuclease (D1740A) domains on a doubly-labelled forked DNA oligonucleotide. Nicking products (indicated with purple triangle) were resolved by denaturing PAGE. **(D)** Cryo-EM density map of the DruE-DruH holocomplex. **(E)** Structural model of the DruE-DruH holocomplex, shown in cartoon representation and in the same orientation as in A. **(F)** Close-up view of the DruE-DruH interaction that is centered on the alpha-helical stalk in the DruE stalk domain. **(G)** Close-up view of the DruE-DruH interaction that is centered on the body of the DruE stalk domain. **(H)** Plaque assays of T2 phage infecting *E. coli* strains expressing wild-type (WT) PaDruantia, PaDruE and PaDruH interface mutants, or a vector-only control. Data represents mean PFU mL^-1^ ± SEM of three independent replicates (n=3).

To obtain structural insights into the molecular architecture of the DruE-DruH complex, we reconstituted DruE and DruH together with a forked DNA substrate (**Figure 4A-B**) and used single-particle cryo-EM analysis to obtain a reconstruction with a nominal resolution of 3.3 Å (**Figure 4D-E & Extended data Fig. 7A-D**). The structure of the DruE-DruH complex reveals that the DruE homodimer disassembles upon interaction with DruH and DNA, forming a 1:1 DruE-DruH heterodimer (**Fig. 4B-C**). DruH contacts the Stalk domain of DruE via its C-terminal alpha helical domain through an extensive interface (∼2130 Å^2^) that involves both electrostatic and hydrophobic interactions (**Figure 4F-G**). Consistent with the structural observations, mutations of the interacting residues in DruE and DruH resulted in a strong reduction of anti-phage activity in *E. coli* (**Figure 4H**).

DruE-DruH complex formation induces large conformational changes in both DruE and DruH, including coordinated domain rearrangements across the RecA, Stalk, Gate and PLD domains of DruE (**Extended data Fig. 8A**). The interaction of DruE with DruH and DNA displaces the Gate domain by ∼13 Å, opening the DNA binding channel and priming of DruE for translocation (**Extended data Fig. 8B**). Concurrently, the PLD domains shift by ∼13 Å to become positioned over the helicase body of the complex (**Extended data Fig. 8C**). This rearrangement is associated with increased flexibility of the PLD domains, as reflected by a decrease in the local resolution (**Extended data Fig. 7C**). Although not directly observed in the cryo-EM map, re-arrangements in the Stalk domain are likely to contribute to the disruption of the DruE dimer. Upon its interaction with DruE and DNA, DruH transitions from a compact to an open conformation, including a displacement of the Ig-1 domain by 58 Å (**Extended data Fig. 8D-F**). Notably, the density corresponding to the DruH N-terminal alpha helical domain is not resolved in the DruE-DruH complex, suggesting that this domain is flexibly associated. Taken together, these data show that DNA substrate-dependent DruE-DruH complex formation is accompanied with extensive molecular rearrangements and reveals critical molecular determinants of the DruE-DruH interaction necessary to support phage defense by type III Druantia.

### DruE preferentially acts on duplex–ssDNA junctions with a 5′ overhang

The structure of the DruE-DruH complex reveals that DruE engages the ssDNA 3’-overhang of a forked DNA substrate (**Figure 4E and 5A**). At the entry site in the helicase core, DruE contacts the backbone of the DNA duplex region via its OB domain (**Figure 4E, 5A & Extended data Fig. 8G**). The ssDNA 3’-overhang is threaded through a positively charged channel towards the RecA-like helicase core, with partially resolved density for nucleobases dT43–dG49 (**Figure 5A & Extended data Fig. 8G**). Nucleobases dG50-dC55 are well resolved and form a continuous stack that is stabilized by contacts with RecA1, whereas RecA2 positioned at the adjacent unresolved ssDNA region (**Figure 5A & Extended data Fig. 8G**). The DNA binding channel is constricted by Asp654^DruE^ that contacts dG50. The downstream nucleotide (dT51) is stabilized by Arg657^DruE^, while dT53 makes a π-π stacking interaction with Phe1292^DruE^ (**Figure 5A & Extended data Fig. 8G**). In agreements with these structural observations, alanine substitution of these residues abolished phage defense by type III Druantia *in vivo* (**Figure 5B**).

**Figure 5:**
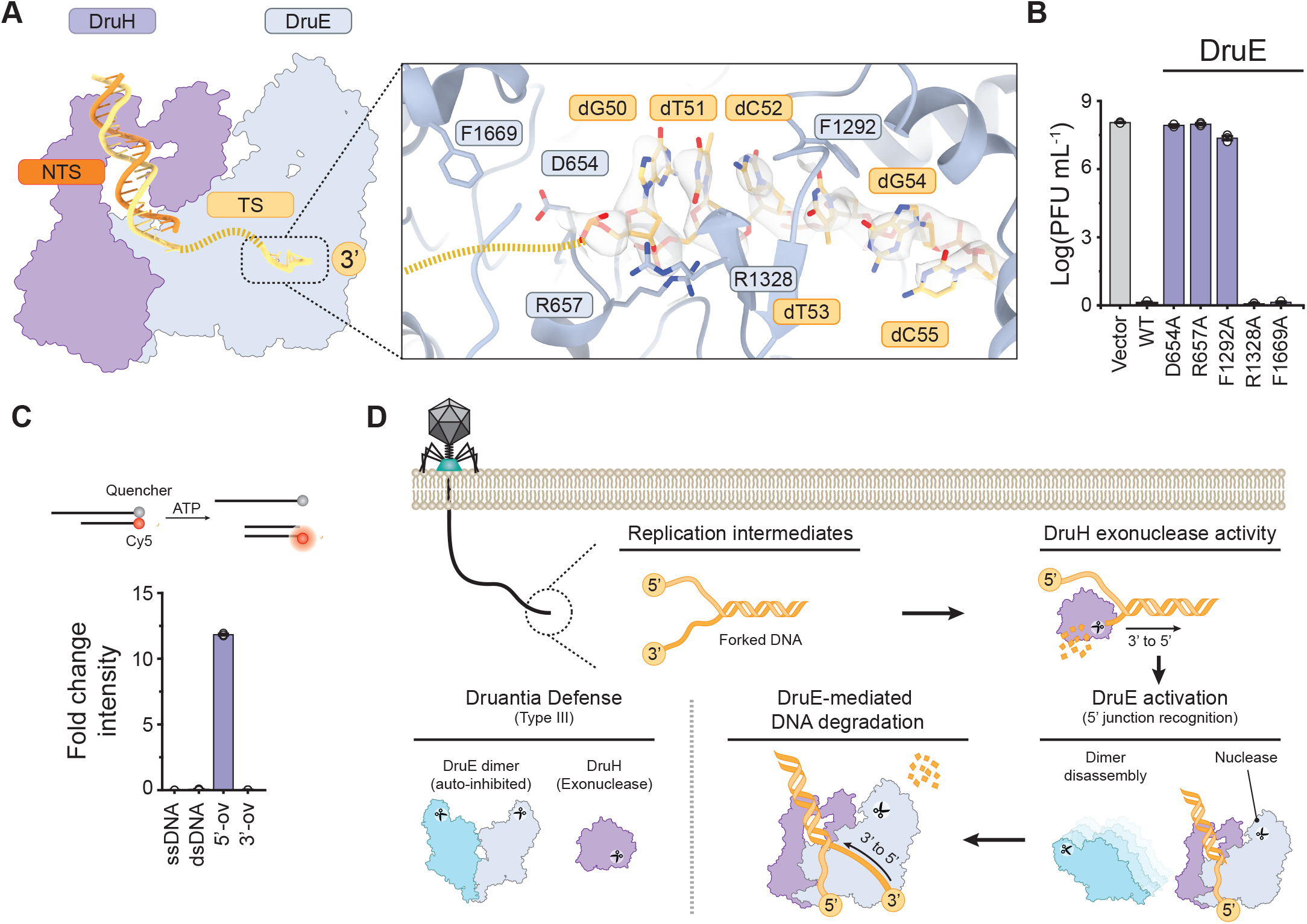
Duplex–ssDNA junction recognition by DruE and mechanistic model for type III Druantia immunity. **(A)** Schematic representation of the DruE-DruH holo complex (right). Inset shows zoomed-in view of DruE interactions with the structurally ordered part of the 3’ ssDNA overhang. **(B)** Plaque assays of T2 phage infecting *E. coli* strains expressing wild-type (WT) PaDruantia, PaDruE helicase mutants, or a vector-only control. Data represents mean PFU mL^-1^ ± SEM of three independent replicates (n=3). **(C)** Top: Schematic showing a fluorophore-quencher assay to probe DNA unwinding by DruE using ssDNA, or dsDNA substrates with blunt ends, a 5’ overhang (5’-ov) or a 3’ overhang (3’-ov). Bottom: Helicase activity of DruE, quantified by change in fluorescence intensity upon incubation of the DNA substrate with DruE and ATP. **(D)** Bacteriophage infection generates replication intermediates containing forked DNA structures. These substrates are initially recognized by DruH, which engages the DNA and processively degrades the 3′ strand through its exonuclease activity. Formation of a 5′ DNA junction together with DruH binding triggers DruE recruitment and activation. DruE dimer disassembly promotes DNA unwinding in a 3′ to 5′ direction, resulting in processive DNA degradation. Together DruE and DruH form an inducible defense complex that selectively targets phage replication intermediates.

The observed polarity of the bound ssDNA overhang is compatible with 3’-5’ translocation and is consistent with the unwinding mechanism of SF2 helicases such as MfrA^23^. To validate the translocation directionality, we performed DNA unwinding assays using fluorophore-quencher DNA substrates (**Figure 5C**). Unexpectedly, unwinding activity was only observed for DNA duplexes containing a 5’-overhang (**Figure 5C**). We hypothesized that DruE recognizes duplex-ssDNA junctions with a 5′ overhang to initiate translocation on the opposing 3′ strand. To test this model, we used substrates containing a biotin moiety at the junction, enabling streptavidin conjugation to act as a steric roadblock to helicase progression. Streptavidin conjugation at the junction reduced the fluorescence amplitude without altering the overall kinetic profile, consistent with a requirement for junction accessibility during the initiation step rather than the rate-limiting translocation process (**Extended data Fig. 8H**). Together, these structural and biochemical data indicate that DruE functions as a 3’-5’ translocase that requires a 5′ overhang for loading on the translocating stand. This is consistent with the observed preference of DruE-DruH for forked DNA.

## Discussion

Our study reveals the molecular mechanism underpinning phage immunity elicited by type III Druantia systems. We demonstrate that DruH encodes a large multidomain protein that functions as a DNA targeting 3’ to 5’ exonuclease, while the helicase-nuclease DruE initially adopts an auto-inhibited dimeric state, which dissociates upon DNA-induced association with DruH. These structural and biochemical analyses support a mechanistic model in which type III Druantia systems selectively target replication-associated forked DNA intermediates generated during phage infection (**Figure 5D**). DruH processes 3’ termini of forked DNA intermediates to generate substrates containing 5′ overhangs. Subsequently, DruH recruits DruE to the duplex-ssDNA junction, inducing DruE dimer disassembly and activation. This enables DruE to undergo 3’ to 5’ translocation, coupled with degradation of both DNA strands. This mechanism enables Druantia to engage phage-specific DNA structures, while avoiding the circular bacterial chromosome. Moreover, the host chromosome is replicated and repaired within tightly controlled nucleoprotein complexes, which likely limits the formation or exposure of forked DNA intermediates that can be targeted by Druantia. The model is consistent with the absence of Druantia-induced toxicity and is compatible with the recombination-based replication mechanism that T-even phages use to amplify their genome during early infection^33^. Together, these results underscore that the type III Druantia innate immune system functions through molecular pattern recognition by sensing a phage-specific pathogen-associated molecular pattern (PAMP) that is absent from the host.

The structural architecture of DruH shows multiple domains including previously uncharacterized folds. The central part of DruH contains tandem repeats of Ig-like folds that resemble adhesion molecules ICAM and NCAM. In eukaryotes, ICAM family proteins mediate cell–cell adhesion and play key roles in immune processes including leukocyte trafficking and immune synapse formation, while NCAM contributes to cell recognition in both neural and immune contexts^34–36^. Although these parallels do not imply functional conservation, they suggest that Ig-like folds represent an evolutionarily versatile structural scaffold that can be repurposed for immune-related functions across domains of life. Structural analysis of the DruH-DruE complex reveals that the C-terminal α-helical domain mediates interaction with DruE, whereas the N-terminal domain does not participate in complex formation. Based on AlphaFold predictions and mutational analysis, we speculate that the N-terminal domain contributes to the exonuclease activity of DruH. The molecular mechanism and catalytic geometry underlying this activity remain unresolved and will require further structural and biochemical investigation.

The sensing of phage-specific PAMP nucleic acid structures represents a recurring theme in bacterial innate immunity. For example, Shedu has been shown to impair phage proliferation by sensing free DNA ends, resulting in dsDNA nicking at a fixed distance from the 5’ end^4^. Similarly, the Lamassu system was shown to release a nuclease effector upon recognition of DNA ends by its SMC-like receptor^5,6^. Our findings extend this paradigm by demonstrating that type III Druantia senses replication-associated forked DNA structures to achieve self– non-self discrimination. DruE is evolutionarily related to the helicase component of DISARM system that relies on DNA methylation to avoid self-targeting^37^. Consistent with our data, DISARM is activated by DNA substrates containing 5′ single-stranded overhangs^37^, suggesting that Druantia and DISARM employ conserved helicase architectures to detect phage-specific DNA configurations, while coupling substrate recognition to distinct downstream effector mechanisms.

Beyond the evolutionary relationship with DISARM, type III Druantia frequently co-occurs with the type II Zorya defense system in bacterial defense islands and has been reported to act synergistically with Zorya in phage defense^18^. Mechanistic studies of Zorya have shown that it encodes a membrane-bound sensor that triggers nuclease activity upon phage sensing^38,39^. Given the dependence of Druantia on enzymatic processing of DNA substrates, we speculate that Zorya may function upstream of Druantia by generating or exposing DNA structures that are optimal for recognition by the DruE-DruH complex. Further studies using pairwise genetic deletions and biochemical reconstitution will be required to determine the functional interdependence of Zorya and Druantia and to establish whether Zorya-mediated processing directly licenses Druantia activation.

The sensing of nucleic acid structures by Druantia raises parallels with eukaryotic innate immune pathways that sense nucleic geometry as PAMPs. In eukaryotes, the immune sensor ZBP1, detects left-handed Z-DNA and Z-RNA conformations that arise during viral infection and triggers programmed cell death^40–42^. Similarly, the vertebrate cytosolic pattern recognition receptor RIG-I specifically recognizes short dsRNA substrates encompassing 5’-triphosphosphate termini, a hallmark of viral replication intermediates, to initiate type-1 interferon signaling^43–45^. The discovery of a prokaryotic antiviral system that recognizes and targets forked DNA substrates thus further underscores the paradigm that recognition of pathogen-specific nucleic acid structures is a conserved feature of innate immunity across the domains of life.

## Data and code availability

Atomic coordinates and cryo-EM maps for the DruE apo dimer (PDB: 29DU), EMDB: EMD-57108), DruH apo (PDB: 29DR), EMDB: EMD-57103) and DruH-DruE complex (PDB: 29DP), EMDB: EMD-57101) have been deposited in the PDB and EMDB databases.

## Acknowledgements

We are grateful to Marta Sawica, Simona Sorrentino and Piotr Szwedziak from University of Zurich Center for Microscopy and Image Analysis and Felix Weis from the European Molecular Biology Laboratory for technical support with cryo-EM data acquisition. We thank Franklin Nobrega, Nicohlas Taylor, Markus Wahl, Chase Beisel for coordinating publication timelines. We thank all the members of the Jinek and Loeff lab for constructive feedback throughout the project. The work was funded by a European Research Council Consolidator Grant (project no. 820152, CRISPR2.0) to MJ. LL was funded by a Horizon 2020 Marie Skłodowska-Curie Individual Fellowship (project no. 845268, MSOPGDM) and a LUMC Junior Principal Investigator Grant 2025. MJ is International Research Scholar of the Howard Hughes Medical Institute, Vallee Scholar of the Vallee Foundation and member of the Swiss National Competence Center for Research “RNA & Disease”.

## Author Contributions

Conceptualization, L.L. and M.J.; Methodology, L.L.; Investigation, LL. and CC; Writing - Original Draft, L.L.; Writing - Review & Editing, L.L. and M.J.; Funding Acquisition, L.L. and M.J.; Resources, L.L. & M.J.; Supervision, L.L.

## Methods

### Bioinformatic analysis of DruE orthologs

DruE sequences of types I to III used for the bioinformatic analysis were retrieved from the Refseq database though the use of Defensefinder. In addition, type 4 DruE sequences were obtained from PADLOC and used as an input for the basic local alignment search tool of NCBI with a 50% sequence identity cutoff ^19,20,49^. All sequences were combined and aligned using MAFFT (v7.490) ^50^, after which a phylogenetic tree was generated using IQ-tree (v2.1.3) ^51^ with automated model selection ^52^ and visualized using the iTOL webserver.

### Plasmid DNA constructs and site-specific mutants

For heterologous expression in *E. coli*, the DNA sequences of *Pseudomonas aeruginosa* BL12 DruE (Genbank: ERY33374.1) and DruH (Genbank: ERY33375.1) were inserted into the 1B (Addgene: 29653) and 2HR-T (Addgene: 29718) plasmids using ligation-independent cloning (LIC), resulting in constructs that carry an N-terminal hexahistidine or hexahistidine-twin-strep-tactin affinity purification tags, respectively, followed by a tobacco etch virus (TEV) protease cleavage site. Site-specific mutations were introduced by QuickChange mutagenesis or by inverse PCR. Plasmids were purified using the GeneJET plasmid miniprep kit (Thermo Fisher Scientific) and mutations were verified by Sanger and full plasmid sequencing.

### Phage cultivation and plaque assays

Phage experiments were performed as described previously^4^. In brief, experiments with phages T2 (DSM16352), T3 (DSM4621), T5 (DSM16353), T6 (DSM4622) and T7 (DSM4623) were performed in *E. coli* strain DSM613, whereas experiments with Lambda phage (DSM4499) were performed in *E. coli* strain K12 (DSM4230). To generate phage stocks, *E. coli* cells were grown in lysogeny broth (LB) supplemented with 1 mM MgCl_2_, at 37 °C, 220 rpm until reaching an optical density at 600 nm (OD_600_) of 0.3. Cultures were subsequently infected with phage and incubation was continued until collapse of the culture. To clean up the lysates, 1% cholorophorm was added together with 10 µg mL^-1^ DNase 1 (NEB) and 1 µg mL^-1^ RNase A, followed by incubation for 1 hour at 37◦C. After incubation, samples were cleared by centrifugation at 15,000g for 10 minutes, supplemented with NaCl to 0.5M and stored at 4◦C until the titer was determined with a plaque assay.

For plaque assays the respective *E. coli* strain was grown in LB until reaching an OD_600_ of 1.0. Next, 400 cells µL were mixed with 50 µL diluted phage stock and 4 mL lukewarm top agar (LB supplemented with 7 g L^-1^ agar and 15 µM IPTG) and poured onto 9 cm LB agar plate with 0.1% L-arabinose at room temperature. Plates were incubated overnight at 37◦C and the next day plaque forming units (PFU) were counted and used to calculate the viral titer. Stocks were diluted with LB supplemented with 0.5M NaCl and 1 mM MgCl_2_ to 10^8^ PFU mL^-1^ and stored at 4◦C until further use.

### Growth curves and infection dynamics in liquid culture

Growth curves and infection assays were performed as described previously^4^. In brief, to probe growth of *E. coli* in the presence or absence of phage, cells were grown in LB until reaching an OD_600_ of 1.0, after which expression was induced with 15 µM IPTG. Cells were diluted with LB with 15 µM IPTG to an OD_600_ of 0.1 and, if indicated, infected with phage to the desired moiety of infection (MOI). Next, 100 µL of the diluted cultures were transferred to a 96-well plate and sealed with a Breathe-Easy sealing membrane (Z380059, Sigma Aldrich). Growth at 37◦C, 300 rpm was tracked for 10 hours by measuring the OD_600_ at 5-minute intervals, using a Spectrostar plate reader (BMG labtech). For each measurement, two technical replicates were averaged prior to determining the average and deviations from the three independent biological replicates.

### Isolation of genomic bacteriophage DNA

Genomic phage DNA was isolated using the DNeasy Blood and Tissue Kit (Qiagen) with the following adjustments. First, phages were precipitated by incubating 10 mL lysate with 8% PEG6000 for 18 hours at 4◦C. Precipitated phages were centrifuged for 30 minutes at 10,000g at 4 °C, after which the supernatant was removed and the pellet was dissolved in 20 mM Tris-HCl (8.0), 250 mM NaCl to a total volume of 200 µL. Next, 20 µL of proteinase K and 200 µl AL Buffer was added and incubated for 20 minutes at 65◦C. After this step, the protocol of the DNeasy Blood and Tissue Kit was followed starting from the addition of 200 µL absolute ethanol.

### Expression and purification of DruE

Hexahistidine-tagged DruE proteins were expressed in *E. coli* BL21-star cells. Cultures were grown at 37 °C (130 rpm) until they reached an OD_600_ of 0.6, after which the cultures were incubated on ice for 1 hour. Protein expression was then induced with 0.25 mM isopropyl-β-D-thiogalactopyranoside (IPTG) and continued for 16 h at 18 °C. Cells were harvested by centrifugation and resuspended in buffer A (20 mM Tris-HCl pH 8.0, 500 mM NaCl, 5 mM imidazole, 1 µg mL^−1^ pepstatin, 200 µg mL^−1^ AEBSF), followed by lysis in a Maximator cell homogenizer at 1,500 bar and 4 °C. The lysate was cleared by centrifugation at 10,000g for 30 min at 4 °C and applied to 15 mL pre-equilibrated Ni-NTA column (Qiagen). The Ni-NTA column was washed with 150 mL of buffer B (20 mM Tris-HCl pH 8.0, 500 mM NaCl, 10 mM imidazole). Proteins were eluted in five fractions (15 mL each) of buffer C (20 mM Tris-HCl pH 8.0, 500 mM NaCl, 250 mM imidazole). Protein-containing elution fractions were pooled and dialyzed overnight against buffer D (20 mM Tris-HCl pH 8.0, 150 mM NaCl, 1 mM DTT) in the presence of TEV protease. To remove uncleaved DruE proteins, the dialyzed fraction was supplemented with 10 mM imidazole and ran over 7.5-mL equilibrated Ni-NTA beads (Qiagen). The flow-through fraction was collected and concentrated using 100 kDa molecular weight cut-off centrifugal filters (Merck Millipore) and the protein was further purified by size-exclusion chromatography using a Superdex 200 (16/600) column (Cytiva) equilibrated in buffer D. Purified proteins were concentrated to 11.5 mg mL^−1^, flash frozen in liquid nitrogen and stored at −80 °C until further use.

### Expression and purification of DruH and StrepII-DruH

Hexahistidine-tagged and Hexahisti-dine-StepII-tagged DruH proteins were expressed in *E. coli* BL21-star cells. Cultures were grown at 37 °C (130 rpm) until they reached an OD_600_ of 0.6, after which the cultures were incubated on ice for 1 hour. Protein expression was then induced with 0.25 mM isopropyl-β-D-thiogalactopyranoside (IPTG) and continued for 16 h at 18 °C. Cells were harvested by centrifugation and resuspended in buffer A (20 mM Tris-HCl pH 8.0, 500 mM NaCl, 5 mM imidazole, 1 µg mL^−1^ pepstatin, 200 µg mL^−1^ AEBSF), followed by lysis in a Maximator cell homogenizer at 1,500 bar and 4 °C. The lysate was cleared by centrifugation at 10,000g for 30 min at 4 °C and applied to 15 mL pre-equilibrated Ni-NTA column (Qiagen). The Ni-NTA column was washed with 150 mL of buffer B (20 mM Tris-HCl pH 8.0, 500 mM NaCl, 10 mM imidazole). Proteins were eluted in five fractions (15 mL each) of buffer C (20 mM Tris-HCl pH 8.0, 500 mM NaCl, 250 mM imidazole). Protein-containing elution fractions were pooled and dialyzed overnight against buffer D (20 mM Tris-HCl pH 8.0, 150 mM NaCl, 1 mM DTT) in the presence of TEV protease. For proteins containing a Strep-tactin affinity tag TEV protease was omitted from the dialysis. To remove uncleaved DruH proteins, the dialyzed fraction was supplemented with 10 mM Imidazole and ran over 7.5-mL equilibrated Ni-NTA beads (Qiagen). The flow-through fraction was collected and and concentrated using 100 kDa molecular weight cut-off centrifugal filters (Merck Millipore) and the protein was further purified by size-exclusion chromatography using a Superdex 200 (16/600) column (Cytiva) equilibrated in buffer D. Purified proteins were concentrated to 6.5 mg mL^−1^, flash frozen in liquid nitrogen and stored at −80 °C until further use.

### DruE-DruH complex reconstitution by affinity co-precipitation

To reconstitute the DruE-DruH complex, 150 pmol StrepII-DruH and 375 pmol DruE were mixed with 225 pmol synthetic DNA substrate in a buffer containing 20 mM Tris-HCl pH 8.0, 50 mM NaCl, and 0.01% Tween20. After 1 hour of incubation at 4◦C, 50 µL of equilibrated Strep-tactin beads (IBA-life sciences) were added to the sample and incubated for 30 minutes at 4◦C on a rotating wheel. Unbound proteins and DNA were removed by washing the beads were 3 times with a buffer containing 20 mM Tris-HCl pH 8.0, 50 mM NaCl. Subsequently, the DruE-DruH complexes were eluted by incubating the beads for 5 minutes at 4◦C with elution buffer containing 20 mM Tris-HCl pH 8.0, 50 mM NaCl and 2.5 mM desthiobiotin.

### *In vitro* DruH nuclease activity assays with fluorescently labelled oligo nucleotides

dsDNA targets were generated by annealing synthetic oligonucleotides (**Extended data Table 2**, IDT) in a buffer containing 10 mM Tris-HCl, 50 mM NaCl using a thermocycler (Biorad) and stored at -20◦C until further use. For the cleavage assays, 40 nM of DNA or RNA target was mixed with 400 nM of DruH in a buffer containing 20 mM Tris-HCl, 100 mM NaCl and 5 mM MgCl_2_ and incubated for the indicated time points at 37◦C. After the incubation, the reaction was quenched by the addition of 1 µL proteinase K and incubated for 10 minutes at 60◦C. Next, samples were mixed with loading dye (97% formamide, 25 mM EDTA and 0.15% Orange G), incubated for 10 minutes at 95◦C and loaded onto a 15% or 10% denaturing PAGE gel containing 8M urea. Gels were run for 2.5 hours at 350 V, followed by imaging with the Typhoon trio (GE healthcare).

### *In vitro* DruE nuclease activity assays with fluorescently labelled oligo nucleotides

For the cleavage assays, 40 nM of DNA target was mixed with 400 nM of DruE in a buffer containing 20 mM Tris-HCl, 100 mM NaCl, 5 mM MgCl2 and 1 mM ATP and incubated for the indicated time points at 37◦C. After the incubation, the reaction was quenched by the addition of 1 µL proteinase K and incubated for 10 minutes at 60◦C. Next, samples were mixed with loading dye (97% formamide, 25 mM EDTA and 0.15% Orange G), incubated for 10 minutes at 95◦C and loaded onto a 15% or 10% denaturing PAGE gel containing 8M urea. Gels were run for 2.5 hours at 350 V, followed by imaging with the Typhoon trio (GE healthcare). All sequences of oligonucleotides used in this study are provided in **Extended data Table 2**.

### *In vitro* cleavage assays with DruE-DruH on genomic phage DNA

For the cleavage assays with genomic DNA, 20 pM of genomic DNA was mixed with 800 pM of DruE and 800 pM DruH in a buffer containing 20 mM Tris-HCl, 50 mM NaCl, 5 mM MgCl_2_, 5% glycerol, 1 mM ATP and incubated for the indicated times at 37◦C. After the incubation, the reaction was quenched by the addition of 1 µL proteinase K. Next, samples were mixed with 6x DNA loading dye (10 mM Tris-HCl (pH 7.6), 60 mM EDTA, 60% Glycerol, 0.03% Bromophenol blue and 0.03% Xylene cyanol FF) and loaded onto a 1% agarose gel. Gels were run for 40 minutes at 100 V, followed by imaging with an ChemiDoc Imaging System (Biorad).

### *In vitro* DruE-DruH DNA degradation assays with fluorescently labelled oligo nucleotides

dsDNA targets were generated by annealing synthetic oligonucleotides (**Extended data Table 2**, IDT) in a buffer containing 10 mM Tris-HCl, 50 mM NaCl using a thermocycler (Biorad) and stored at -20◦C until further use. For DruE-DruH DNA degradation assays, 40 nM of DNA target was mixed with 400 nM of DruH and 1200 nM DruE in a buffer containing 20 mM Tris-HCl, 50 mM NaCl, 5 mM MgCl2, 5% Glycerol and 1 mM ATP and incubated for the indicated time points at 37◦C. After the incubation, the reaction was quenched by the addition of 1 µL proteinase K and incubated for 10 minutes at 60◦C. Next, samples were mixed with loading dye (97% formamide, 25 mM EDTA and 0.15% Orange G), incubated for 10 minutes at 95◦C and loaded onto a 15% or 10% denaturing PAGE gel containing 8M urea. Gels were run for 2.5 hours at 350 V, followed by imaging with the Typhoon trio (GE healthcare).

### *In vitro* DruE DNA unwinding assays

dsDNA targets were generated by annealing synthetic oligonucleotides (**Extended data Table 2**, IDT) in a buffer containing 10 mM Tris-HCl, 50 mM NaCl using a thermocycler (Biorad) and stored at -20◦C until further use. For DNA unwinding assays, 50 nM dsDNA target was mixed with 250 nM DruE in a buffer containing 20 mM Tris-HCl, 50 mM NaCl, 5 mM MgCl_2_, 5% glycerol, and 0.05% Tween20. After incubation for 15 minutes at 4◦C, samples were transferred to a 96-well plate and 1 mM ATP together with 500 nM unlabeled competitor DNA oligonucleotide was added. The samples were incubated for 5 min at room temperature pior to measuring the fluorescence intensity using a PHERAstar FSX Microplate Reader (BMG Labtech).

For kinetic measurements of DNA unwinding, 50 nM dsDNA target was mixed with 1 mM ATP, 500 nM Streptavidin and 100 nM unlabeled competitor DNA oligonucleotide in a buffer containing 20 mM Tris-HCl, 50 mM NaCl, 5 mM MgCl_2_, 5% glycerol, and 0.05% Tween20. After incubation for 45 minutes at 4◦C, samples were transferred to a 96-well plate and 50 nM DruE was added. The fluorescence intensity was measured immediately over 20 minutes the using a CLARIOstar Microplate Reader (BMG Labtech).

### *In vitro* ATPase assays

The ATPase activity of WT and mutant DruE on single-stranded and double stranded DNA substrates (**Extended data Table 2**) was measured using the EnzCheck Phosphate kit (Invitrogen) at 26°C. Here, the provided 2-amino-6-mercapto-7-methylpurine riboside (MESG) substrate together with the released inorganic phosphate (P_i_) is enzymatically converted to 2-amino-6-mercapto-7-methyl-purine which absorbs light at 360 nm, thus, facilitating the measurement of the P_i_ concentration in the reaction mixture. For ATPase activity assays, 200 nM DruE was incubated with 20 nM DNA substrate in a buffer containing 20 mM Tris-HCl, 250 mM NaCl and 5 mM MgCl_2_, in a total volume of 200 µL. The ATPase reactions were initiated by adding 0.5 mM ATP to the samples. Absorbance at 360 nm was measured once per minute over 45 min in a PHERAstar FSX Microplate Reader (BMG Labtech).

### Sample preparation and cryo-EM data collection of apo DruH

For cryo-EM grid preparation, 300-mesh Amorphous Nickel-Titanium Holey Foil Grids (Au R1.2/1.3, ANTcryo™) was glow-discharged for 60 seconds at 15 mA (PELCO easiGlow™, Ted Pella). 2.5 µL DruH sample at 1.5 mg mL^-1^ was applied, and the grids were blotted for 6 s at 4 °C using and 80% humidity using 50 mm Vitrobot filter paper (Electron Microscopy Sciences). Grids were plunge-frozen in liquid ethane using a Vitrobot Mark IV plunger (Thermo Fisher Scientific) and stored in liquid nitrogen until cryo-EM data collection. Cryo-EM data collection was performed on a Titan Krios G3i microscope (Thermo Fisher Scientific) equipped with a Gatan K3 direct electron detector operated at 300 kV in super-resolution counting mode. Data acquisition was performed using the EPU automated data acquisition software with three shots per hole at defocus range of −0.6 μm to −2.0 μm (0.2-µm steps). The final dataset comprised a total of 9,516 micrographs at a calibrated magnification of 130,000x and a super-resolution pixel size of 0.325 Å. Micrographs were exposed for 1.26 s with a total dose of 66.21 e^−^ Å^−2^ over 47 subframes.

### Data processing and model building of apo DruH

Cryo-EM data was processed using cryoSPARC v4.4.1 ^53^. Imported micrographs were motion-corrected with patch motion correction (multi) and CTF values were estimated using patch CTF estimation (multi). An initial set of particles was picked with blob picker using a circular blob and a minimum and maximum particle diameter of 90 and 130 Å, respectively. After extraction of the particles with a box size of 400×400 pixels, particles were subjected to 2D classification with a Maximum resolution of 12 Å, an initial classification uncertainty factor of 1 and with a circular mask of 110 Å to generate templates for picking (5 templates).

After template-based picking with a particle diameter of 130 Å, classes with defined particles were selected, resulting in a total of 1,500,264 particles, which were used to generate two *ab initio* models of which one was used for heterogeneous refinement with three classes. Classes were inspected visually using UCSF Chimera^54^, and the particles and volume of the best class were subjected to a second round of *ab initio* model generation with four classes and the class similarity setting set to 0.9. The particles and volume of the best class were used as an input for per-particle motion correction and subsequently refined using non-uniform refinement with optimization of CTF parameters and per-particle defocus. The final map was sharpened with a B-factor of -137. The local resolution was estimated based on the resulting map using the local resolution function of cryoSPARC and plotted on the map using UCSF Chimera^54^. The structural model of Apo DruH was built in Coot (V0.9.2) ^55^ and was refined over multiple rounds using Phenix ^56,57^. Real-space refinement was performed with the global minimization, atomic displacement parameter (ADP) refinement and secondary structure restrains enabled. The quality of the atomic model, including protein geometry, Ramachandran plots, clash analysis and model cross-validation, was assessed with MolProbity and validation tools in Phenix ^56–59^. The refinement statistics of the final model are listed in **Extended data Table 1**. Figures of maps, models and the calculations of map contour levels were generated using ChimeraX^54^.

### Sample preparation and cryo-EM data collection of apo DruE

For cryo-EM grid preparation, 300-mesh holey carbon grids (Au R1.2/1.3, Quantifoil Micro Tools) was glow-discharged for 60 seconds at 15 mA (PELCO easiGlow™, Ted Pella). 2.5 µL DruE sample at 2.5 mg mL^-1^ was applied, and the grids were blotted for 4 s at 4 °C using and 80% humidity using 50 mm Vitrobot filter paper (Electron Microscopy Sciences). Grids were plunge-frozen in liquid ethane using a Vitrobot Mark IV plunger (Thermo Fisher Scientific) and stored in liquid nitrogen until cryo-EM data collection. Cryo-EM data collection was performed on a Titan Krios G3i microscope (Thermo Fisher Scientific) equipped with a Gatan K3 direct electron detector operated at 300 kV in super-resolution counting mode. Data acquisition was performed using the EPU automated data acquisition software with three shots per hole at defocus range of −1.0 μm to −2.4 μm (0.2-µm steps). The final dataset comprised a total of 9,689 micrographs at a calibrated magnification of 130,000x and a super-resolution pixel size of 0.325 Å. Micrographs were exposed for 1.26 s with a total dose of 59.35 e^−^ Å^−2^ over 47 subframes.

### Data processing and model building of apo DruE

Cryo-EM data was processed using cryoSPARC v4.4.1^53^. Imported micrographs were motion-corrected with patch motion correction (multi) and CTF values were estimated using patch CTF estimation (multi). An initial set of particles was picked with blob picker using a circular blob and a minimum and maximum particle diameter of 170 and 220 Å, respectively. After extraction of the particles with a box size of 480×480 pixels, particles were subjected to 2D classification with a circular mask of 220 Å. Classes with defined particles were selected, resulting in a total of 1,303,729 particles, which were used to generate two *ab initio* models of which one was used for heterogeneous refinement with four classes. Classes were inspected visually using UCSF Chimera^54^, and the particles and volume of the best class were subjected to a second round of *ab initio* model generation with four classes and the class similarity setting set to 0.9. The particles and volume of the best class were used as an input for per-particle motion correction and subsequently refined using non-uniform refinement with optimization of CTF parameters and per particle defocus, after which refinement was repeated with relaxed-C2 symmetry enabled. The final map was sharpened with a B-factor of -113. The local resolution was estimated based on the resulting map using the local resolution function of cryoSPARC and plotted on the map using UCSF Chimera^54^. The structural model of Apo DruE was built in Coot (V0.9.2) ^55^ and was refined over multiple rounds using Phenix ^56,57^. Real-space refinement was performed with the global minimization, atomic displacement parameter (ADP) refinement and secondary structure restrains enabled. The quality of the atomic model, including protein geometry, Ramachandran plots, clash analysis and model cross-validation, was assessed with MolProbity and validation tools in Phenix^56–59^. To aid interpretation, the map was feature enhanced with DeepEMhancer^60^. The refinement statistics of the final model are listed in **Extended data Table 1**. Figures of maps, models and the calculations of map contour levels were generated using ChimeraX^54^.

### Sample preparation and cryo-EM data collection of DruE-DruH holo complex

Prior to grid preparation for cryo-EM, 1.5 µM StrepII-DruH was combined with 3.75 µM DruE and 2.5 µM forked DNA substrate in a buffer containing 20 mM Tris-HCl pH 8.0, 50 mM NaCl, and 0.01% Tween20, after which 50 µL of equilibrated Strep-tactin beads (IBA-life sciences) were added to the sample and incubated for 45 minutes at 4◦C on a rotating wheel. Unbound proteins and DNA were removed by washing the beads were 3 times with a buffer containing 20 mM Tris-HCl pH 8.0, 50 mM NaCl, and 0.01% Tween20 and DruE-DruH complexes were eluted by incubating the beads for 5 minutes at 4◦C with elution buffer containing 20 mM Tris-HCl pH 8.0, 50 mM NaCl, and 2.5 mM desthiobiotin. Eluted proteins were concentrated to 1.0 mg mL^-1^ using 100 kDa molecular weight cut-off centrifugal filters (Merck Millipore). For cryo-EM grid preparation, 300-mesh holey carbon grids (Au R1.2/1.3, Quantifoil Micro Tools) were glow-discharged for 60 seconds at 15 mA (PELCO easiGlow™, Ted Pella). 2.5 µL samples were applied and blotted for 4 s at 80% humidity and 4 °C using 50 mm Vitrobot filter paper (Electron Microscopy Sciences). Grids were plunge frozen in liquid ethane using a Vitrobot Mark IV plunger (Thermo Fisher Scientific) and stored in liquid nitrogen until cryo-EM data collection. Cryo-EM data collection was performed on a Titan Krios G3i microscope (Thermo Fisher Scientific) equipped with a Gatan K3 direct electron detector operated at 300 kV in super-resolution counting mode. Data acquisition was performed using the EPU automated data acquisition software with three shots per hole at defocus range of −1.0 μm to −2.4 μm (0.2-µm steps). The final dataset comprised a total of 7,773 micrographs at a calibrated magnification of 130,000x and a super-resolution pixel size of 0.325 Å. Micrographs were exposed for 1.26 s with a total dose of 62.93 e^−^ Å^−2^ over 37 subframes.

### Data processing and model building of DruE-DruH holo complex

Cryo-EM data was processed using cryoSPARC (v4.7.1)^53^. Imported micrographs were motion-corrected with patch motion correction (multi) and CTF values were estimated using patch CTF estimation (multi). An initial set of particles was picked with blob picker using a circular blob and a minimum and maximum particle diameter of 80 and 170 Å, respectively. After extraction of the particles with a box size of 420×420 pixels, particles were subjected to 2D classification to generate templates for picking (5 templates). After template-based picking with a particle diameter of 140 Å, particle picks were cleaned up with micrograph junk detector, extracted and subjected to 2D classification with a circular mask of 180 Å. Classes with defined particles were selected, resulting in a total of 529,118 particles, which were used to generate two *ab initio* models of which one was used for heterogeneous refinement with three classes. Classes were inspected visually using UCSF Chimera^54^, and the particles and volume of the best class were used as an input heterogeneous refinement with two classes. The best class was used for 3D classification, after which one class was used for per-particle motion correction and subsequent refinement using non-uniform refinement with optimization of per-particle defocus and CTF parameters. The final map was sharpened with a B-factor of -54. The local resolution was estimated based on the resulting map using the local resolution function of cryoSPARC and plotted on the map using UCSF Chimera ^54^. The structural model of the DruE-DruH was built in Coot ^55^ and refined over multiple rounds using Phenix ^56,57^. Real-space refinement was performed with global minimization, atomic displacement parameter (ADP) refinement and secondary structure restrains enabled. The quality of the atomic model, including protein geometry, Ramachandran plots, clash analysis and model cross-validation, was assessed with MolProbity and the validation tools in Phenix^56–59^. To aid interpretation, the map was feature enhanced with DeepEMhancer^60^. The refinement statistics of the final model are listed in **Extended data Table 1**. Figures of maps, models and the calculations of map contour levels were generated using ChimeraX ^54^.

