## Supplementary information for "Processing of forked DNA activates a helicase-nuclease immune system"

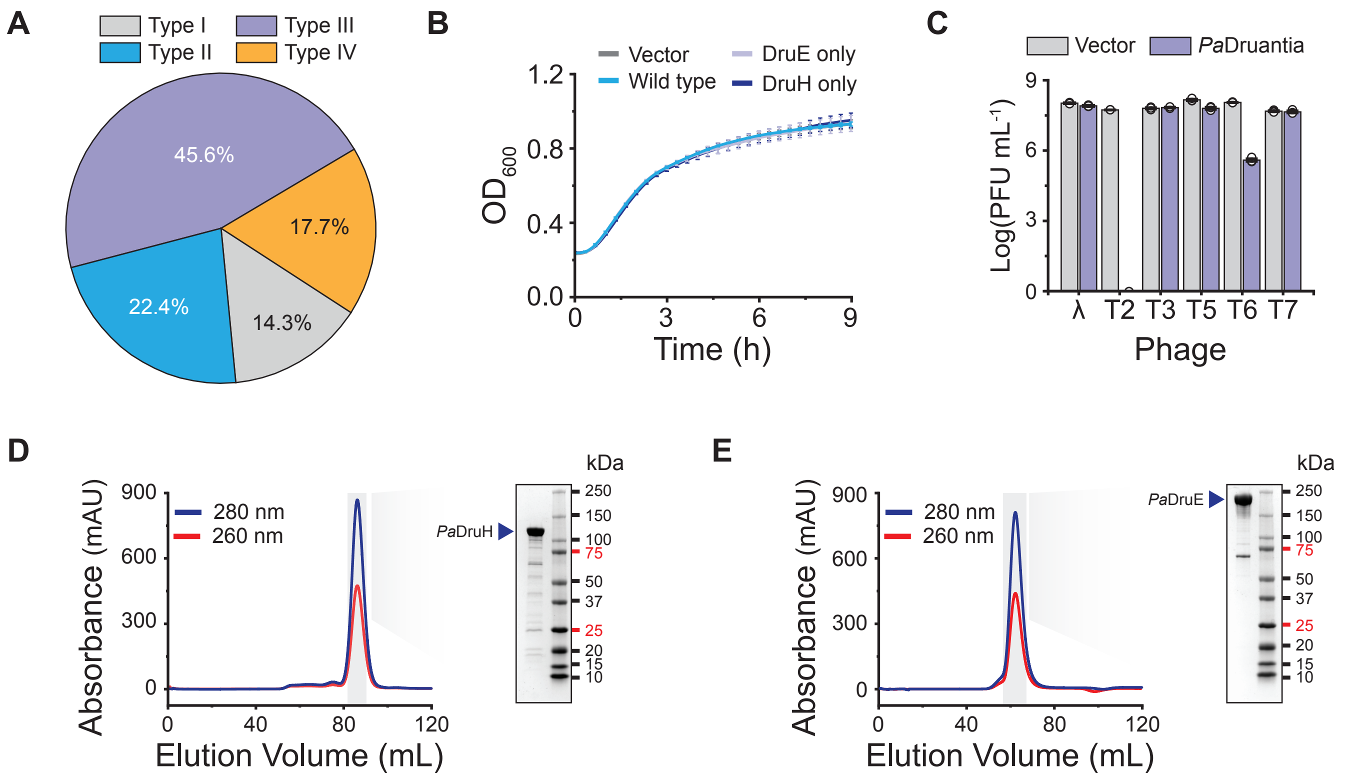


**Extended Data Fig. 1: Bioinformatic analysis, phage assays and purification of Druantia phage defense systems.**

**(A)** Distribution of Druantia types. **(B)** Growth curves of *E. coli* cells expressing *P. aeruginosa* Druantia, DruH and DruE. Negative control strain harbors empty expression vector. Data points represent mean ± standard error of the mean (SEM) of three biological replicates (n=3). **(C)** Plaque assays of *E. coli* strains expressing type III Druantia using a panel of phages. Data represents mean PFU mL$^{-1}$ ± SEM of three independent replicates (n=3). **(D)** Analysis of purified recombinantly-expressed PaDruH by size-exclusion chromatography (left) and SDS-PAGE (right). **(E)** Analysis of purified recombinantly-expressed PaDruE by size-exclusion chromatography (left) and SDS-PAGE (right).

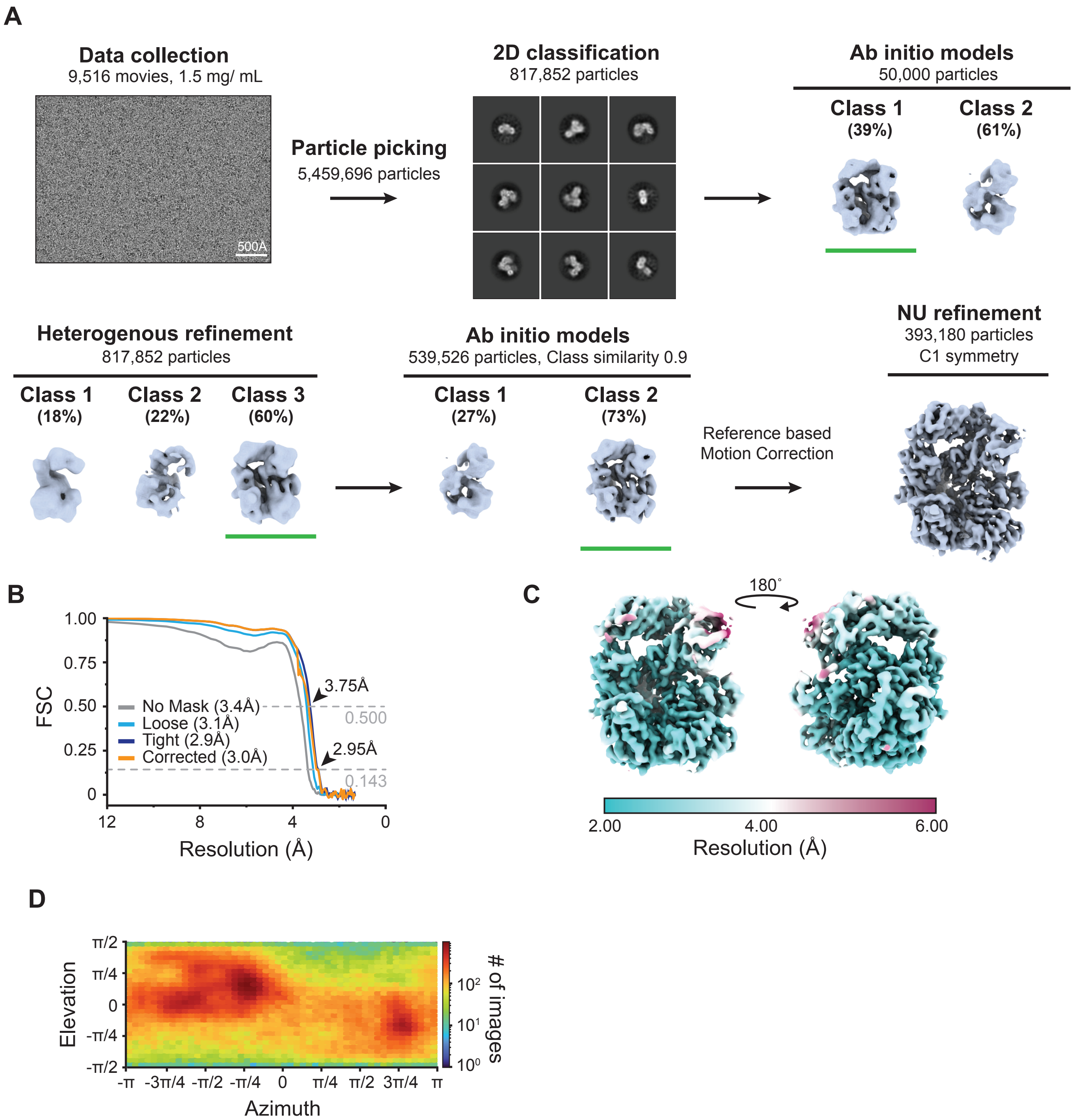


**Extended Data Fig. 2: Cryo-EM processing workflow for the DruH protein.**

**(A)** Cryo-EM processing workflow for DruH. **(B)** FSC determined from two independently refined half-maps. The gold standard cut-off (FSC=0.143) is marked with an arrow. **(C)** Local resolution estimation on the final cryo-EM density map of DruH. **(D)** Euler diagram showing orientation distribution of final cryo-EM reconstruction.

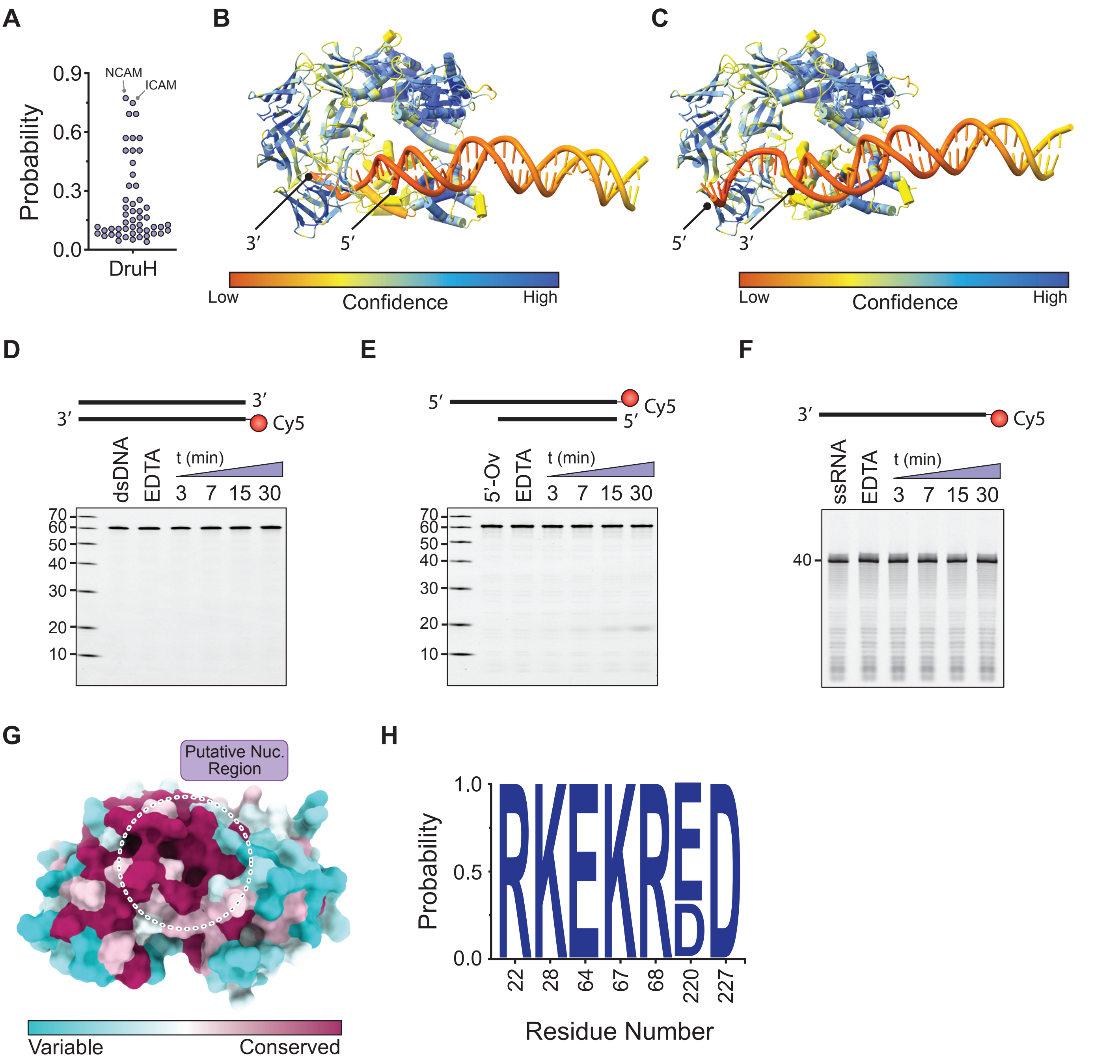


**Extended Data Fig. 3: Molecular details of the DruH protein.**

**(A)** Probability scores from a structural homology search using Foldseek with DruH as query, each circle represents an individual hit[46]. **(B)** Alphafold3 structure prediction of DruH in complex with a partial DNA duplex containing a 3'-ssDNA overhang. **(C)** Alphafold3 structure prediction of DruH in complex with a partial DNA duplex containing a 5'-ssDNA overhang. **(D)** *In vitro* nuclease activity assays with DruH using fluorescently labelled dsDNA oligonucleotides. Cleavage products were resolved by denaturing polyacrylamide gel electrophoresis (PAGE). **(E)** *In vitro* nuclease activity assays with DruH using fluorescently labelled oligonucleotides containing 5'-overhangs. Cleavage products were resolved by denaturing polyacrylamide gel electrophoresis (PAGE). **(F)** *In vitro* nuclease activity assays with DruH using fluorescently labelled ssRNA oligonucleotides. Cleavage products were resolved by denaturing polyacrylamide gel electrophoresis (PAGE). **(G)** Side view of the DruH N-terminal domain colored by sequence conservation. The putative nuclease region is indicated by a dotted circle. **(H)** Sequence conservation analysis of the putative active site residues in DruH. Figure was generated using WebLogo[47].

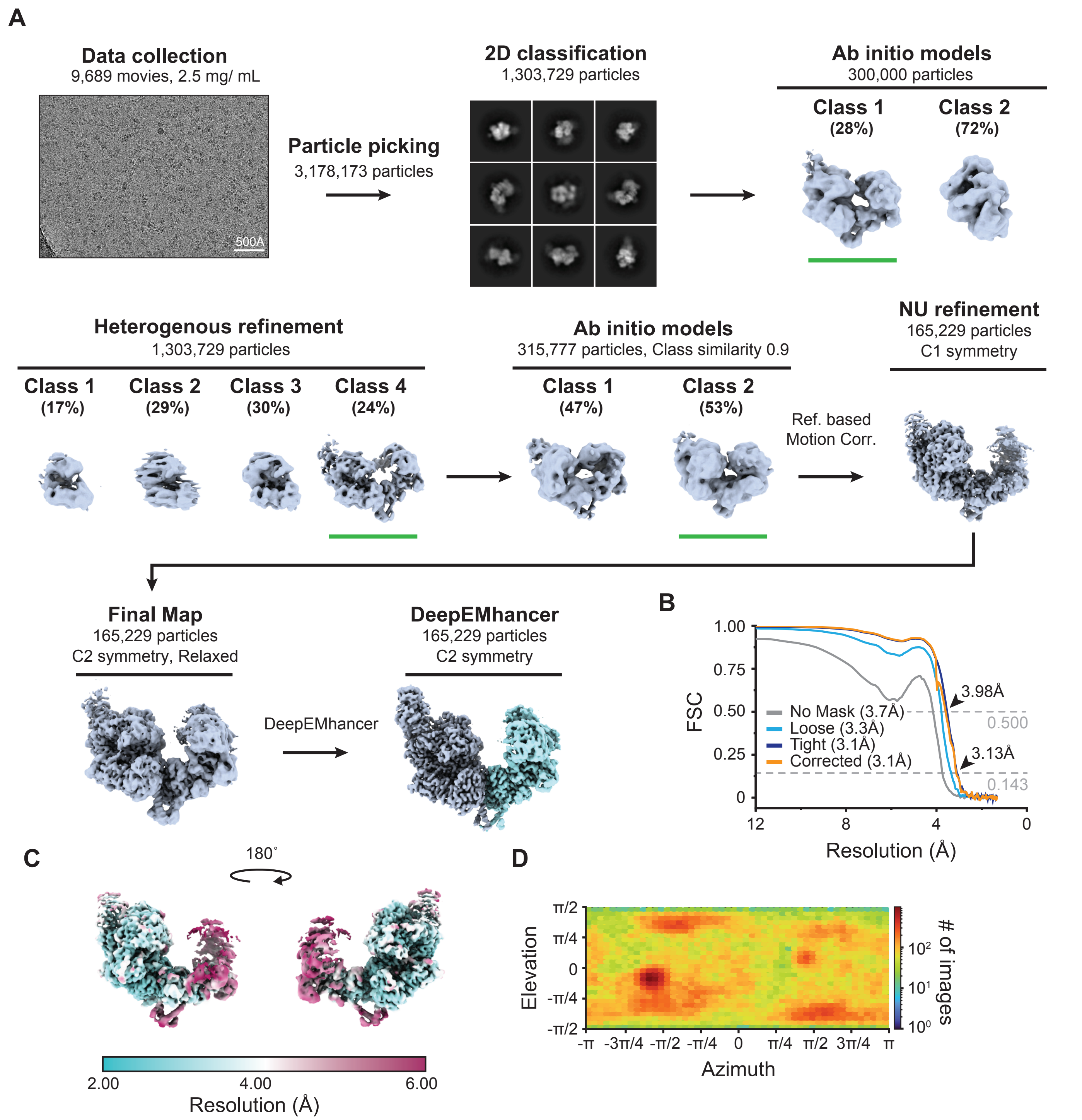


**Extended Data Fig. 4: Cryo-EM processing workflow for the DruE homodimer complex.**

**(A)** Cryo-EM processing workflow for the DruE homodimer complex. **(B)** FSC determined from two independently refined half-maps. The gold standard cut-off (FSC=0.143) is marked with an arrow. **(C)** Local resolution estimation on the final cryo-EM density map of the DruE complex. **(D)** Euler diagram showing orientation distribution of final cryo-EM reconstruction.

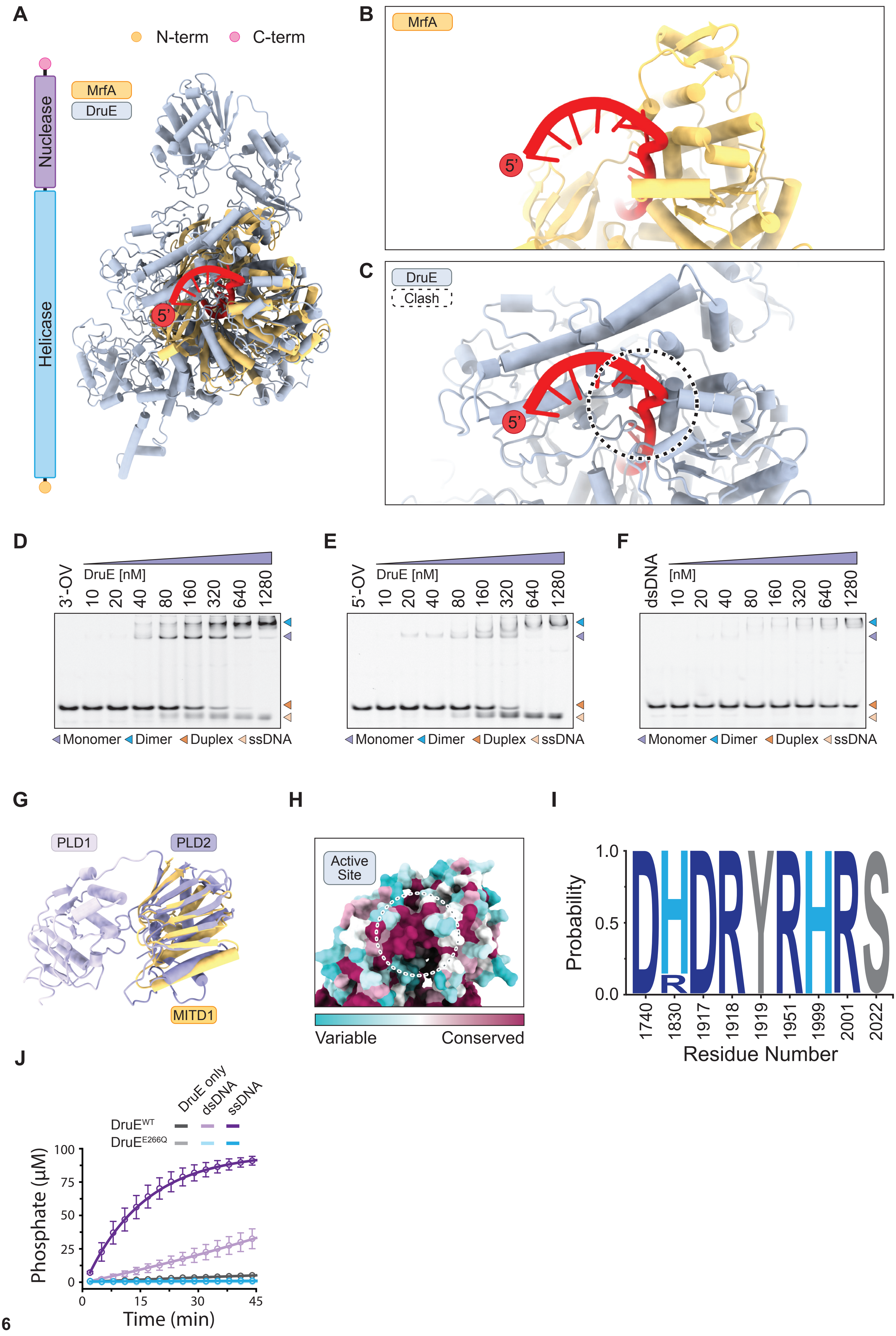

A
N-term
C-term
Nuclease
Helicase
MrfA
DruE
5'
B
MrfA
5'
C
DruE
Clash
5'
D
3'-OV
DruE [nM]
10 20 40 80 160 320 640 1280
Monomer Dimer Duplex ssDNA
E
5'-OV
DruE [nM]
10 20 40 80 160 320 640 1280
Monomer Dimer Duplex ssDNA
F
dsDNA
[nM]
10 20 40 80 160 320 640 1280
Monomer Dimer Duplex ssDNA
G
PLD1
PLD2
MITD1
H
Active Site
Variable
Conserved
I
Probability
1.0
0.5
0.0
1740 1830 1917 1918 1919 1951 1999 2001 2022
Residue Number
J
DruE only
dsDNA
ssDNA
DruE^WT
DruE^E266Q
Phosphate (µM)
100 75 50 25 0
0 15 30 45
Time (min)

**Extended Data Fig. 5: Molecular details of the DruE homodimer complex.**

**(A)** Structural superposition of DruE (grey) with MfrA bound to single-stranded DNA (MfrA orange, DNA red, PDB: 6ZNP)[23]. **(B)** Close-up view of the MfrA (orange) bound to DNA (red, PDB: 6FWS). **(C)** Close-up view of DruE (grey) superimposed with ssDNA as bound in the structure of DNA-bound MfrA (PDB: 6FWS). Structural clashes are indicated by a dotted circle. **(D)** Electrophoretic mobility shift analysis of DruE binding to a fluorescently labelled partial duplex containing a ssDNA 3-overhang. **(E)** Electrophoretic mobility shift analysis of DruE binding to a fluorescently labelled partial duplex containing a ssDNA 5-overhang. **(F)** Electrophoretic mobility shift analysis of DruE binding to a fluorescently labelled dsDNA duplex. **(G)** Structural superposition of DruE (purple) with MITD1 (orange, PDB: (2YMB)[48]. **(H)** Zoom in view of the DruE C-terminal PLD nuclease domain colored by sequence conservation. The nuclease active site is indicated by a dotted circle. **(I)** Sequence conservation analysis of the active site residues in DruE. Figure was generated using WebLogo[47]. **(J)** ATPase activity of DruE variants on ssDNA and dsDNA substrates.

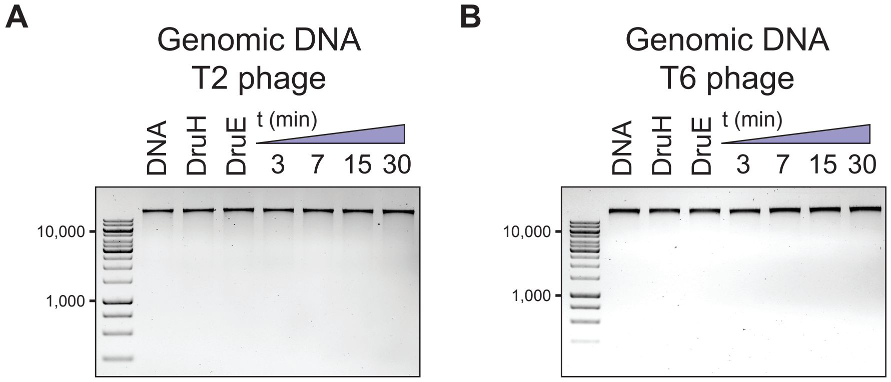


**Extended Data Fig. 6: Nuclease activity of Type III Druantia on genomic DNA.**

**(A)** Analysis of nuclease activity of Type III Druantia on genomic DNA isolated from T2 phages. T2 and T6 extensively modify their DNA by glucosylation[31,32]. **(B)** Analysis of nuclease activity of Type III Druantia on genomic DNA isolated from T2 phages. T2 and T6 extensively modify their DNA by glucosylation[31,32].

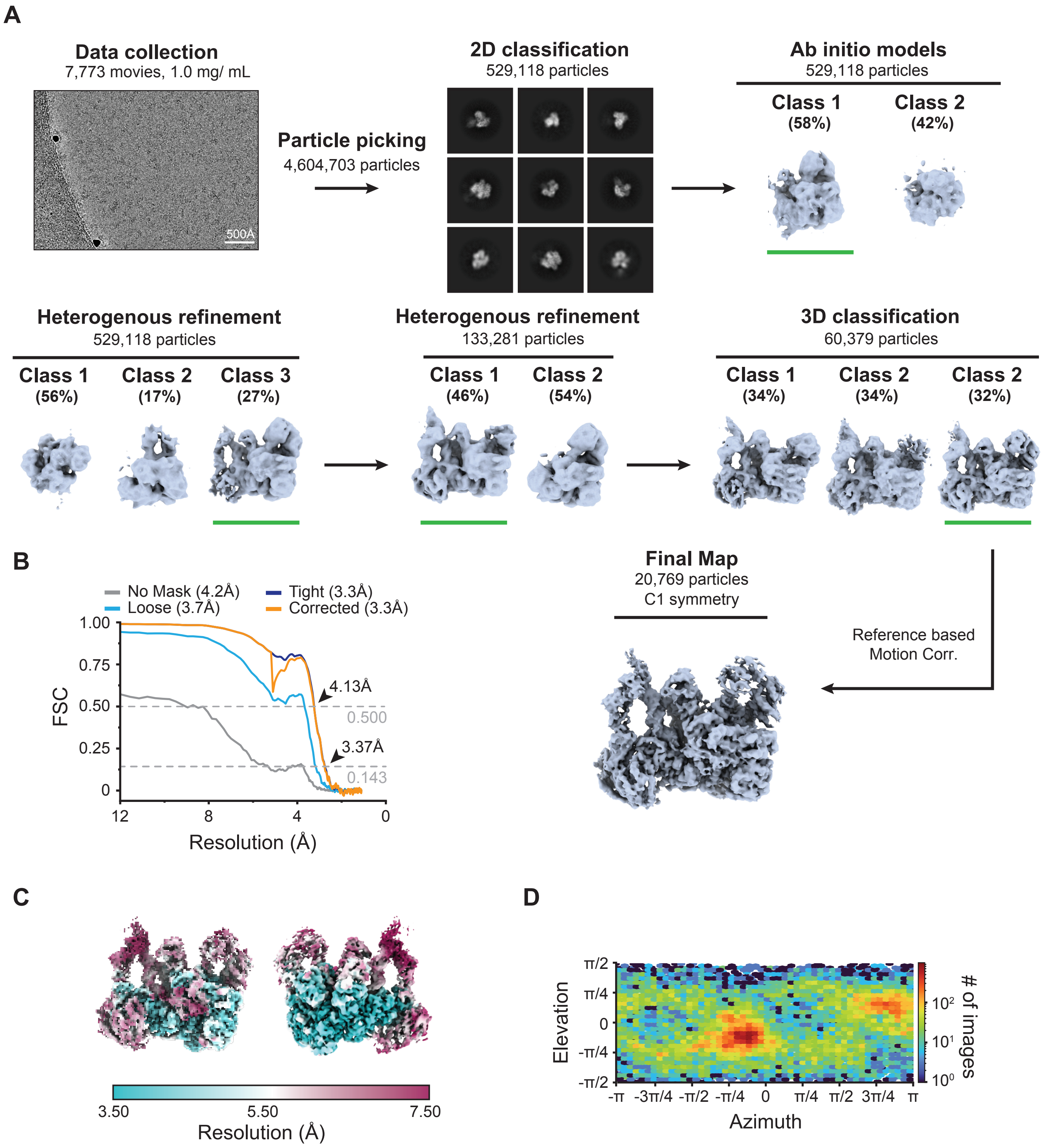


**Extended Data Fig. 7: Cryo-EM processing workflow for the DruE-DruH holo complex.**

(A) Cryo-EM processing workflow for the DruE-DruH holo complex. (B) FSC determined from two independently refined half-maps. The gold standard cut-off (FSC=0.143) is marked with an arrow. (C) Local resolution estimation on the final cryo-EM density map of the DruE-DruH holo complex. (D) Euler diagram showing orientation distribution of final cryo-EM reconstruction.

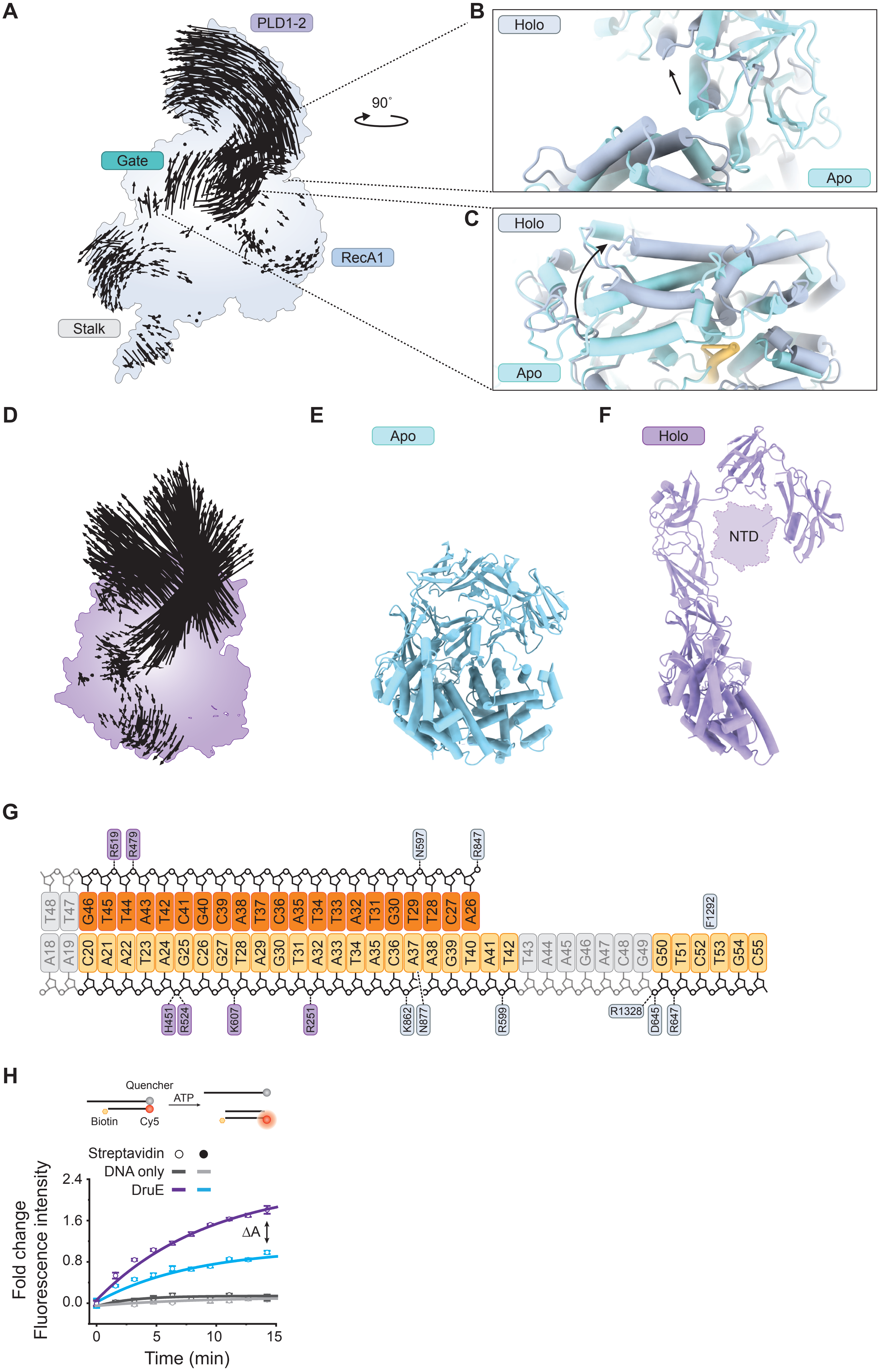
A
PLD1-2
Gate
RecA1
Stalk
90°
B
Holo
Apo
C
Holo
Apo
D
E
Apo
F
Holo
NTD
G
R519
R479
N597
R847
T48 T47 G46 T45 T44 A43 T42 C41 G40 C39 A38 T37 C36 A35 T34 T33 A32 T31 G30 T29 T28 C27 A26
A18 A19 C20 A21 A22 T23 A24 G25 C26 G27 T28 A29 G30 T31 A32 A33 T34 A35 C36 A37 A38 G39 T40 A41 T42 T43 A44 A45 G46 A47 C48 G49 G50 T51 C52 T53 G54 C55
F1292
H451
R524
K607
R251
K862
N877
R599
R1328
D645
R647
H
Quencher
ATP
Biotin
Cy5
Streptavidin
DNA only
DruE
2.4
1.6
0.8
0.0
ΔA
Fold change
Fluorescence intensity
0
5
10
15
Time (min)

**Extended Data Fig. 8: Molecular details of the DruE-DruH holo complex.**

**(A)** Vector map displaying the structural rearrangements in DruE upon DNA binding and complex formation with DruH. **(B)** Zoom-in views of the PLD nuclease domain in apo DruE (cyan) and DruE in complex with DruH and forked DNA (grey). **(C)** Zoom-in views of the gate helix in apo DruE (cyan) and DruE in complex with DruH and forked DNA (grey). **(D)** Vector map displaying the structural rearrangements in DruH upon DNA binding and complex formation with DruE. **(E)** Orthogonal view of apo DruH in compacted form. **(F)** Orthogonal view of DruH in complex with DNA and DruE. Notably, the N-terminal domain (NTD) is not visible in the holo complex structure. **(G)** Schematic showing interactions of DruE (grey) and DruH (purple) with the forked DNA substrate. Opaque grey nucleotides are not visible in the DruE-DruH holo cryo-EM density map. **(H)** *In vitro* DNA unwinding assays with a DNA substrate containing a 5'-overhang and a 3'-biotin moiety. Assays were performed in the presence and absence of streptavidin.
